# A learning-to-rank framework for large-scale identification of unknown proteins from cryo-EM and cryo-ET maps

**DOI:** 10.64898/2026.10.12.758677

**Authors:** Qi Hu, Georges Mer

## Abstract

Cryo-electron microscopy (cryo-EM) and cryo-electron tomography (cryo-ET) can yield density maps whose molecular identities are unknown. Predicted-structure databases enable proteome-scale searches, but distinguishing the correct candidates remains challenging, particularly for intermediate-resolution maps without clear side-chain features. We developed CryoAssign, an automated framework that combines large-scale structure matching with learning-to-rank prioritization. CryoAssign searches intact density maps without prior segmentation and integrates complementary measures of map–model agreement. On held-out benchmark maps, CryoAssign robustly prioritized the correct candidates. It also identified proteins or narrowed the set of candidates in previously unassigned cryo-EM maps and recovered multiple components from complex cryo-EM and cryo-ET maps. Controlled low-pass filtering showed generally robust recovery across a broad range of intermediate resolutions. CryoAssign provides a scalable approach for identifying proteins from cryo-EM and cryo-ET density maps.

---

Recent advances in cryo-electron microscopy (cryo-EM), including computational developments in image preprocessing, three-dimensional reconstruction and classification^1–3^, have enabled structure determination of many biological systems that were previously difficult or impossible to study^4,5^. Cryo-EM is especially powerful for samples available in limited quantities or exhibiting pronounced conformational or compositional heterogeneity^6,7^, allowing high-resolution structural characterization of macromolecular complexes in multiple states and of multiple distinct complexes from a single dataset.

More recently, developments in *in situ* cryo-electron tomography (cryo-ET), particularly with cryo-focused ion beam (cryo-FIB) milling, have enabled visualization of cellular and molecular structures in their native environments with unprecedented detail^8^. Because these approaches are used to image complex native samples, density maps corresponding to unknown molecular components are often obtained.

Several computational approaches have been developed to identify unknown proteins from cryo-EM maps. The advent of AlphaFold2^9,10^ has greatly expanded the availability of high-quality predicted structures and the number of candidate models that can be searched against experimental density maps. Deep learning-based tools such as ModelAngelo^11^ and CryoAtom^12^ perform particularly well for high-resolution maps, typically better than approximately 4 Å, where amino acid side-chain density is sufficiently resolved.

The Situs package^13^ includes CoLoRes^14^, a rigid-body docking program based on exhaustive six-dimensional search and Fourier correlation theory. CoLoRes has been widely used to dock atomic models into electron-density maps and low-resolution envelopes across a broad range of resolutions.

Despite these advances, identifying proteins from intermediate-resolution cryo-EM maps remains challenging because reliable side-chain information is often unavailable and candidate structures must be distinguished from thousands of alternatives. Here, we present CryoAssign, an automated pipeline for large-scale identification of unknown proteins from cryo-EM density maps. The workflow integrates CoLoRes rigid-body docking with automated preprocessing, postprocessing and learning-to-rank prioritization. Candidate structures, including AlphaFold2-predicted models and experimentally determined structures from the Protein Data Bank (PDB), are prepared and docked into target maps. Features describing map–model agreement, structural continuity and docking quality are extracted from the resulting poses and used to prioritize likely candidates. We applied CryoAssign to several previously unexplained cryo-EM maps and complex cryo-ET datasets.

## Results

### CryoAssign automates docking of AlphaFold2 models into cryo-EM maps

To identify unknown proteins from cryo-EM maps at scale, we developed CryoAssign, which automates CoLoRes rigid-body docking of candidate structures. Although CoLoRes has previously been used for protein identification^15^, CryoAssign adds automated preprocessing, parallel screening and candidate reranking, making searches of tens of thousands of structures practical.

CryoAssign can use either AlphaFold2-predicted models or experimentally determined structures in the PDB as input candidates. In a typical workflow, a candidate structure database is first assembled from the AlphaFold2 database, the PDB, or a user-supplied collection of coordinate files (**Fig. 1**). For AlphaFold2 models, additional preprocessing is often required because the relative orientations of domains in full-length predicted models may be unreliable. To address this issue, CryoAssign uses a strategy similar to DomainFit^16^ and applies phenix.process_predicted_model^17,18^ to split multidomain AlphaFold2 models into individual domains based on confidence measures such as predicted local distance difference test (pLDDT) and predicted aligned error (PAE).

**Fig. 1.**
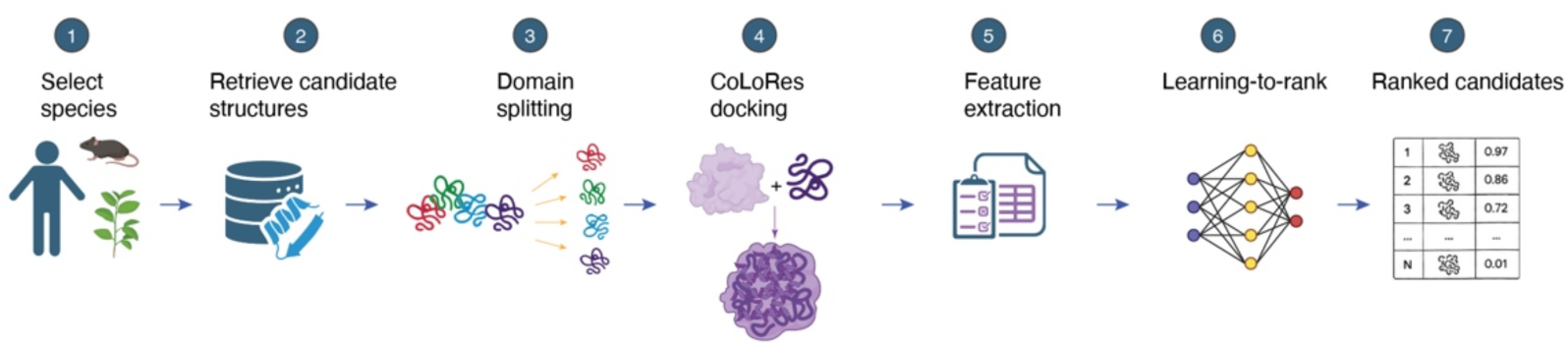
CryoAssign workflow for identifying unknown proteins from cryo-EM density maps. Protein structures are retrieved from species-specific AlphaFold2 or PDB databases. AlphaFold2 models are optionally split into individual domains using phenix.process_predicted_model, whereas experimentally determined PDB structures are used directly following redundancy reduction. Candidate structures are docked independently into the target density map using CoLoRes. Docking-derived features are extracted and integrated by a trained learning-to-rank model to generate a prioritized list of candidate proteins for downstream validation. Created with BioRender.com.

In a typical species-specific search, CryoAssign downloads the corresponding AlphaFold2 models, splits them into domains and docks the resulting domains in parallel against the intact target map using CoLoRes exhaustive six-dimensional search. Unlike DomainFit^16^, CryoAssign does not require prior map segmentation. Docking results are postprocessed and prioritized by an XGBoost-based learning-to-rank model^19,20^ using derived map–model agreement features and CoLoRes quantities.

CryoAssign can also use coordinate files downloaded from the PDB. Because the PDB contains many redundant entries, a prefiltered nonredundant structure database is preferable and can significantly reduce computational cost. When searching against a PDB-derived database, the domain-splitting step is often omitted since oligomeric and multichain assemblies may themselves provide informative docking candidates.

### A learning-to-rank framework improves candidate prioritization

CoLoRes typically generates up to ten poses per candidate and ranks them by correlation-based scores. These scores identify the best pose for a given candidate but compare candidates poorly because they depend on protein size, overlap volume and map resolution. Consequently, raw CoLoRes cross-correlation does not reliably rank thousands of candidates.

Correct fits also show spatial features not captured by the raw score. We therefore derived Pearson correlation- and continuity-based metrics, including correlations within strict and locally dilated overlap masks, and trained a learning-to-rank model to integrate them.

Benchmarking on 14 held-out maps containing at least one manually confirmed true-positive candidate showed that the engineered docking-derived features captured complementary aspects of map–model agreement (**Fig. 2** and **Supplementary Fig. 1**). Although individual features frequently outperformed the original CoLoRes cross-correlation score for subsets of the benchmark, no single feature consistently achieved the best overall ranking performance (**Fig. 2d,g**). Because split AlphaFold2 domains can yield multiple true hits for a map, candidate prioritization is a ranking rather than binary classification problem. We therefore trained an XGBoost-based learning-to-rank model to integrate complementary features using manually annotated training data. Performance was evaluated using Recall@P, map-wise average precision (AP) and normalized discounted cumulative gain (NDCG) (Methods), which quantify how effectively true hits are concentrated near the top of the ranked list. On the held-out maps, the ranker outperformed the original CoLoRes score and the individual engineered features (**Fig.** 2c,e–g,k).

**Fig. 2.**
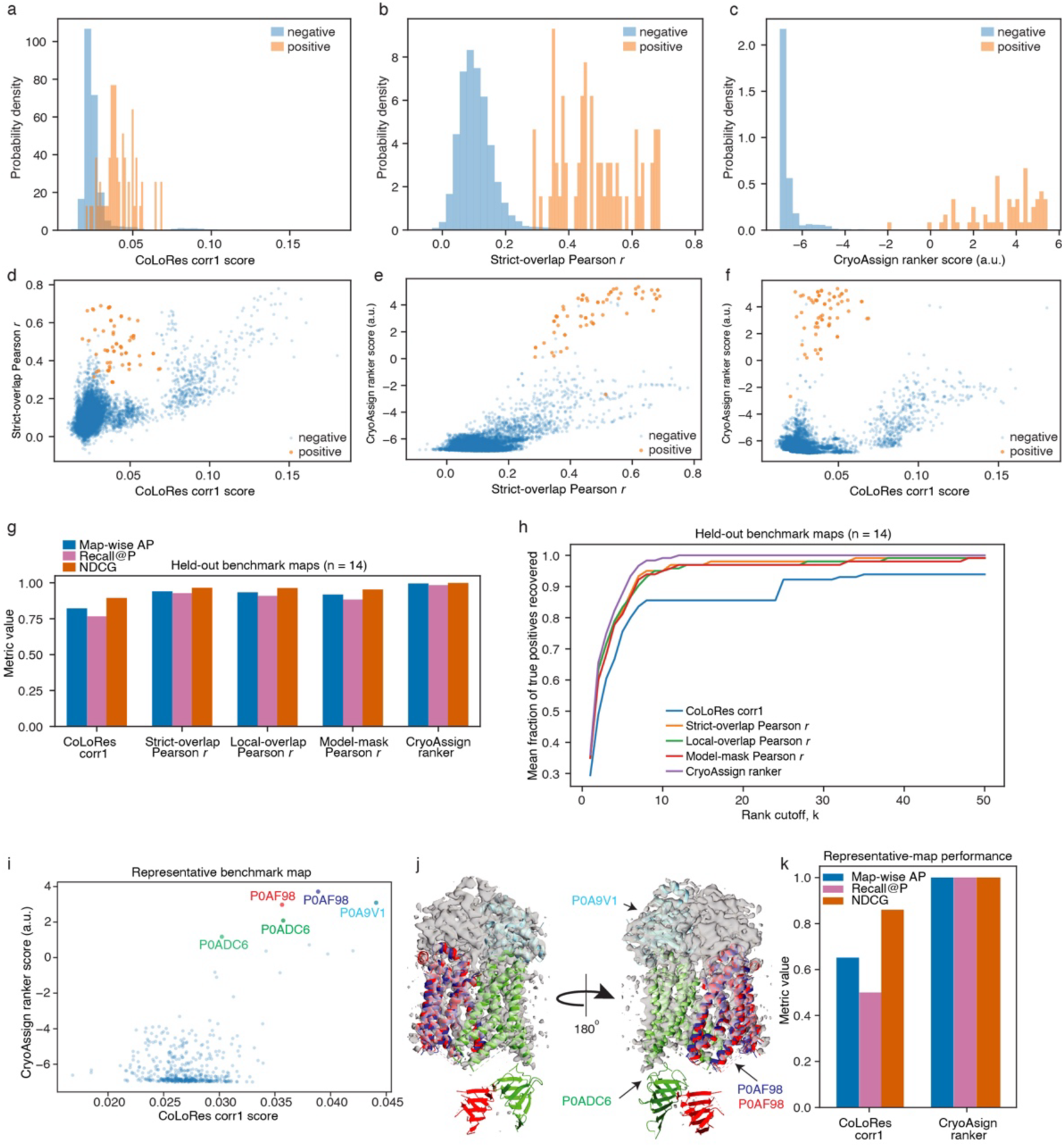
Learning-to-rank integration of complementary docking-derived features improves candidate prioritization. Representative results from the 14 held-out benchmark maps used for recovery-oriented evaluation. **a–c,** Distributions of the CoLoRes corr1 score, Strict-overlap Pearson r, and CryoAssign ranker score, respectively, for positive and negative candidates. CryoAssign ranker scores are shown in arbitrary units (a.u.). **d–f,** Pairwise relationships between (**d**) CoLoRes corr1 score and Strict-overlap Pearson *r*, (**e**) Strict-overlap Pearson *r* and CryoAssign ranker score, and (**f**) CoLoRes corr1 score and CryoAssign ranker score. **g,** Ranking performance on the 14 held-out benchmark maps using the CoLoRes corr1 score, Strict-overlap Pearson *r*, Local-overlap Pearson *r*, Model-mask Pearson *r*, or the CryoAssign ranker. Performance was evaluated using map-wise average precision (Map-wise AP), recall at P (Recall@P), and normalized discounted cumulative gain (NDCG). **h,** Cumulative recovery curves showing the mean fraction of true positives recovered across the held-out benchmark maps as a function of rank cutoff, k, for the same five ranking criteria. **i–k,** Representative results for a single benchmark map, including (**i**) a scatter plot of CoLoRes corr1 score versus CryoAssign ranker score, (**j**) the corresponding cryo-EM density overlaid with correctly identified proteins, and (**k**) per-map ranking performance comparing the CoLoRes corr1 score and CryoAssign ranker. In **g** and **k**, blue, purple, and vermilion denote Map-wise AP, Recall@P, and NDCG, respectively.

### CryoAssign identifies DnaK and ArnA in previously unassigned *E. coli* cryo-EM maps

To assess CryoAssign in a practical experimental setting, we analyzed two unassigned cryo-EM reconstructions obtained from a recombinant protein sample purified from *E. coli* (**Fig. 3a,b**). One reconstruction represented a small particle of approximately 50 kDa and was resolved to approximately 4.2 Å, whereas the second represented a D3-symmetric hexameric particle and was resolved to approximately 3.9 Å. Because the only prior information was that both particles were likely *E. coli* contaminants, we searched each map against a candidate database derived from the *E. coli* AlphaFold2 proteome, which contains 4,370 proteins.

**Fig. 3.**
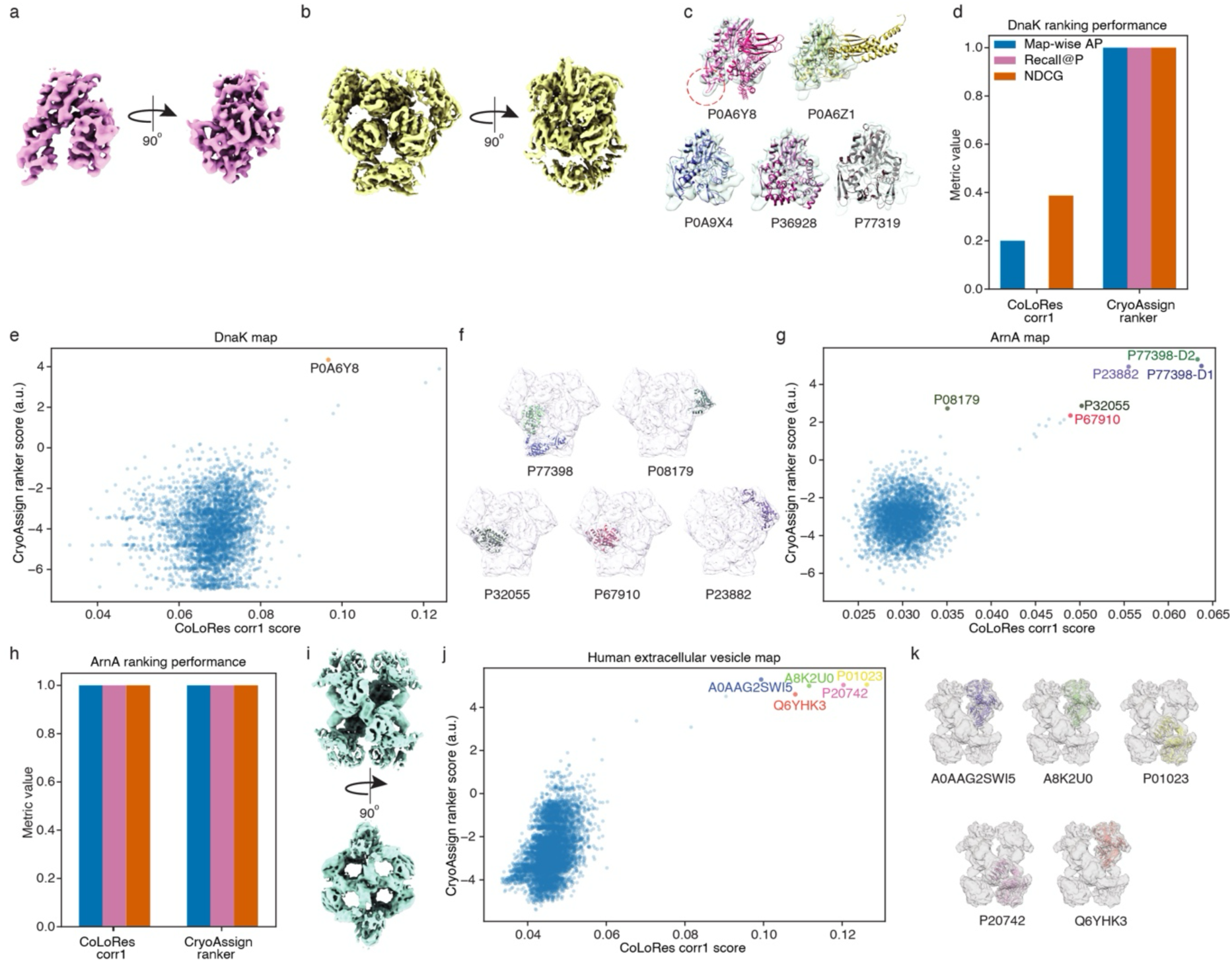
CryoAssign identifies proteins in representative cryo-EM density maps. **a–h,** Identification of the HSP70-family chaperone DnaK and the bifunctional enzyme ArnA. **a,** DnaK density maps in two orthogonal views. **b,** ArnA density maps in two orthogonal views. **c,** DnaK density overlaid with the five highest-ranked candidates; DnaK (P0A6Y8) shows the best agreement. **d,** Ranking performance for the DnaK map using map-wise average precision (map-wise AP), recall at P (Recall@P) and normalized discounted cumulative gain (NDCG). **e,** CoLoRes corr1 score versus CryoAssign ranker score for the DnaK map; DnaK is highlighted. **f,** ArnA density overlaid with the five highest-ranked candidates; ArnA alone accounts for the complete protomer density. **g,** CoLoRes corr1 score versus CryoAssign ranker score for the ArnA map; the correct ArnA domains are highlighted. **h,** Ranking performance for the ArnA map. **i–k,** Identification of human α2-macroglobulin. **i,** Density map in two orthogonal views. **j,** CoLoRes corr1 score versus CryoAssign ranker score; the five highest-ranked candidates belong to the α2-macroglobulin superfamily. **k,** Density overlaid with the five highest-ranked candidates. CryoAssign ranker scores are in arbitrary units. In **d** and **h**, blue, purple and vermilion denote map-wise AP, Recall@P and NDCG, respectively.

For the 4.2-Å reconstruction, the five highest-ranked candidates were the HSP70-family chaperone DnaK (P0A6Y8), chaperone protein HscC (P77319), chaperone protein HscA (P0A6Z1), cell shape-determining protein MreB (P0A9X4) and chaperone protein YegD (P36928) (**Fig. 3c**). Although these candidates have broadly similar overall shapes at the docking resolution, CryoAssign ranked DnaK first (**Fig. 3c–e**). Inspection of the docked models showed that DnaK provided the best agreement with the experimental density, whereas the remaining candidates contained clear local mismatches, supporting assignment of the particle to DnaK.

DnaK is a 638-residue HSP70-family molecular chaperone comprising an N-terminal nucleotide-binding domain and a C-terminal substrate-binding domain^21^. Only the nucleotide-binding domain was resolved in the reconstruction, corresponding to a particle of approximately 40–42 kDa. The resolved conformation was most consistent with an apo or GrpE-bound state, and no density was observed for bound ATP or ADP^22^.

The 3.9-Å reconstruction displayed D3 symmetry, with each protomer containing two distinct domains. SDS–PAGE analysis of the cross-linked sample showed bands at approximately 400 and 70 kDa, consistent with the hexamer and protomer, respectively (**Fig. 3a**). CryoAssign ranked bifunctional polymyxin-resistance protein ArnA (P77398) first, followed by methionyl-tRNA formyltransferase (P23882), GDP-l-fucose synthase (P32055), phosphoribosylglycinamide formyltransferase (P08179) and ADP-l-glycero-d-manno-heptose-6-epimerase (P67910) (**Fig. 3f–h**).

Although all five candidates matched portions of the density, the four alternatives contained fewer than 400 residues and accounted for only part of the protomer density (**Fig. 3f**). ArnA alone accounted for the complete density and had the expected molecular mass of approximately 70 kDa. Moreover, ArnA is known to form a hexamer, consistent with the observed architecture, whereas the other candidates are reported as monomers or homodimers.

ArnA comprises an N-terminal transformylase domain and a C-terminal dehydrogenase domain connected by a short linker^23,24^. Docking the full-length AlphaFold2 protomer produced a slight mismatch between the domains, whereas docking them independently markedly improved agreement with the density, consistent with the relatively high PAE between them. Cryo-EM analysis also revealed conformational heterogeneity between the domains and among subunits of the hexamer (**Supplementary Movie 1**).

### CryoAssign narrows an extracellular vesicle protein map to α2-macroglobulin-family candidates

In another application, an unknown density map was obtained from extracellular vesicles isolated from human tissue culture. The approximately 4.5-Å map exhibited tetrameric D2 symmetry (**Fig. 3i**). Without prior knowledge of its identity, we docked the map against the AlphaFold2 human proteome database, which contains 23,586 proteins.

The five highest-ranked candidates all belonged to the α2-macroglobulin superfamily^25,26^ and shared highly similar domain architectures, making discrimination based solely on local structural features difficult (**Fig. 3j,k**). However, they differed in their oligomeric states. CD109 (Q6YHK3) primarily forms heteromeric complexes^27^, whereas α2-macroglobulin-like protein 1 (A8K2U0) is predominantly monomeric^28^. Pregnancy zone protein (P20742) exists mainly as a dimer, although tetrameric intermediates have been reported^29^. The observed D2-symmetric tetrameric architecture provided an additional structural constraint, narrowing the candidate list to α2-macroglobulin (P01023) and ovostatin homolog 2-like (A0AAG2SWI5). Among these candidates, α2-macroglobulin is the most plausible assignment as it is a well-established extracellular plasma protein that has been repeatedly reported in extracellular vesicle proteomic datasets^30–32^.

### CryoAssign prioritizes multiple proteins from a complex cryo-EM map

To evaluate the ability of CryoAssign to recover multiple correct protein assignments from a single density map, we analyzed the cryo-EM reconstruction EMD-22475 corresponding to the radial spoke complex from the axoneme of *Chlamydomonas reinhardtii*^33^. This dataset represents a particularly challenging test case because the density map contains contributions from numerous protein components with diverse sizes, folds, and levels of local resolution. In contrast to the previous examples, where the objective was to identify a single unknown protein, successful analysis of this map requires simultaneous recovery of many true positives from a large candidate database.

Among the 18 proteins assigned in the corresponding structural model (PDB 7JTK), CryoAssign identified and ranked 11 within the top 15 candidates (**Fig. 4a,b**). By comparison, the original CoLoRes-derived corr1 score ranked only six of these proteins within the top 15 (**Fig. 4b,c**). Two additional proteins, Q39580 and Q27YU0, were ranked 37th and 39th, respectively, by the CryoAssign ranker, whereas corr1 ranked them 143rd and 175th. These results demonstrate that the learning-to-rank framework enriches true positives near the top of the ranked candidate list.

**Fig. 4.**
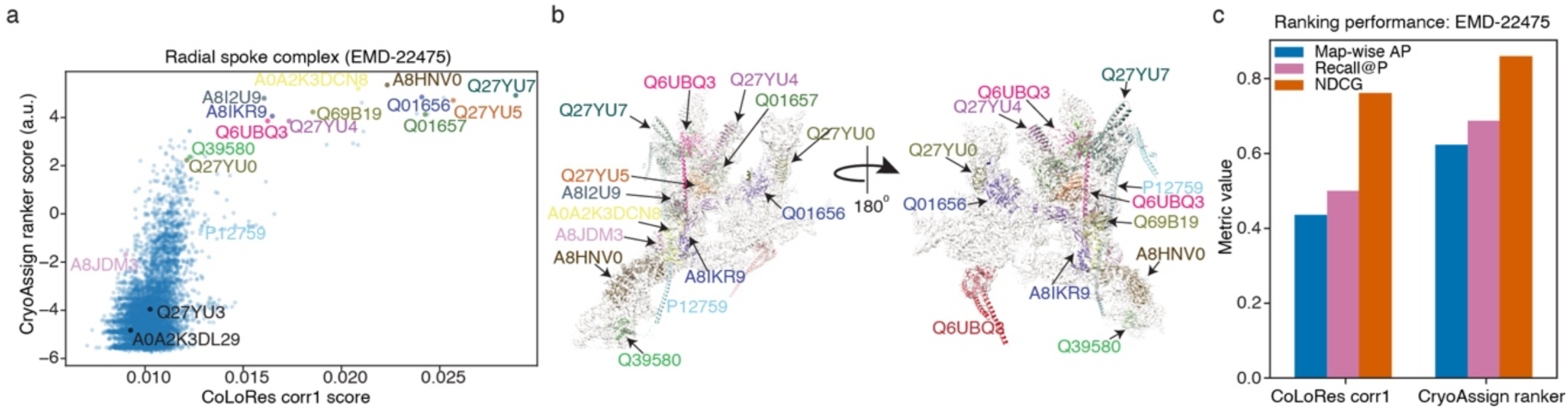
CryoAssign identifies multiple protein components from the cryo-EM map of the radial spoke complex from the *Chlamydomonas reinhardtii* axoneme. **a,** Scatter plot of CoLoRes corr1 score versus CryoAssign ranker score for the radial spoke complex map EMD-22475. Known positive components are labeled. CryoAssign ranker scores are shown in arbitrary units. **b,** Cryo-EM density map of the radial spoke complex overlaid with the correctly identified protein components. Colors correspond to the labeled candidates in panel **a**. **c,** Ranking performance for EMD-22475 comparing the CoLoRes corr1 score and CryoAssign ranker using Map-wise AP, Recall@P, and NDCG. Blue, purple, and vermilion denote Map-wise AP, Recall@P, and NDCG, respectively.

CryoAssign also outperformed the original corr1 score across map-wise average precision, Recall@P and NDCG (**Fig. 4c**). These results support the use of CryoAssign to recover multiple protein assignments from complex cryo-EM and cryo-ET maps.

### Application of CryoAssign to cryo-ET subtomogram-averaged maps

To evaluate the applicability of CryoAssign to cryo-electron tomography, we analyzed a series of subtomogram-averaged (STA) maps spanning resolutions from ∼4 to ∼10 Å. In each case, the maps were searched against the complete species-specific AlphaFold2-derived structure database, providing a stringent test of large-scale protein identification under conditions commonly encountered in in situ structural biology.

One representative example is the recently determined membrane-associated mTORC2 complex (EMD-56117)^34^. The STA map was searched against the complete human AlphaFold2 database comprising 40,596 processed domains. Domain splitting of the four constituent proteins (PDB 9TPW) yielded ten candidate domains. CryoAssign ranked eight of these ten domains within the top ten candidates (**Fig. 5a–c**). The remaining two positions in the top ten (ranks 8 and 10) were occupied by structurally similar proteins. These results demonstrate that CryoAssign effectively prioritizes the correct molecular components from a proteome-scale candidate database, even for challenging cryo-ET reconstructions.

**Fig. 5.**
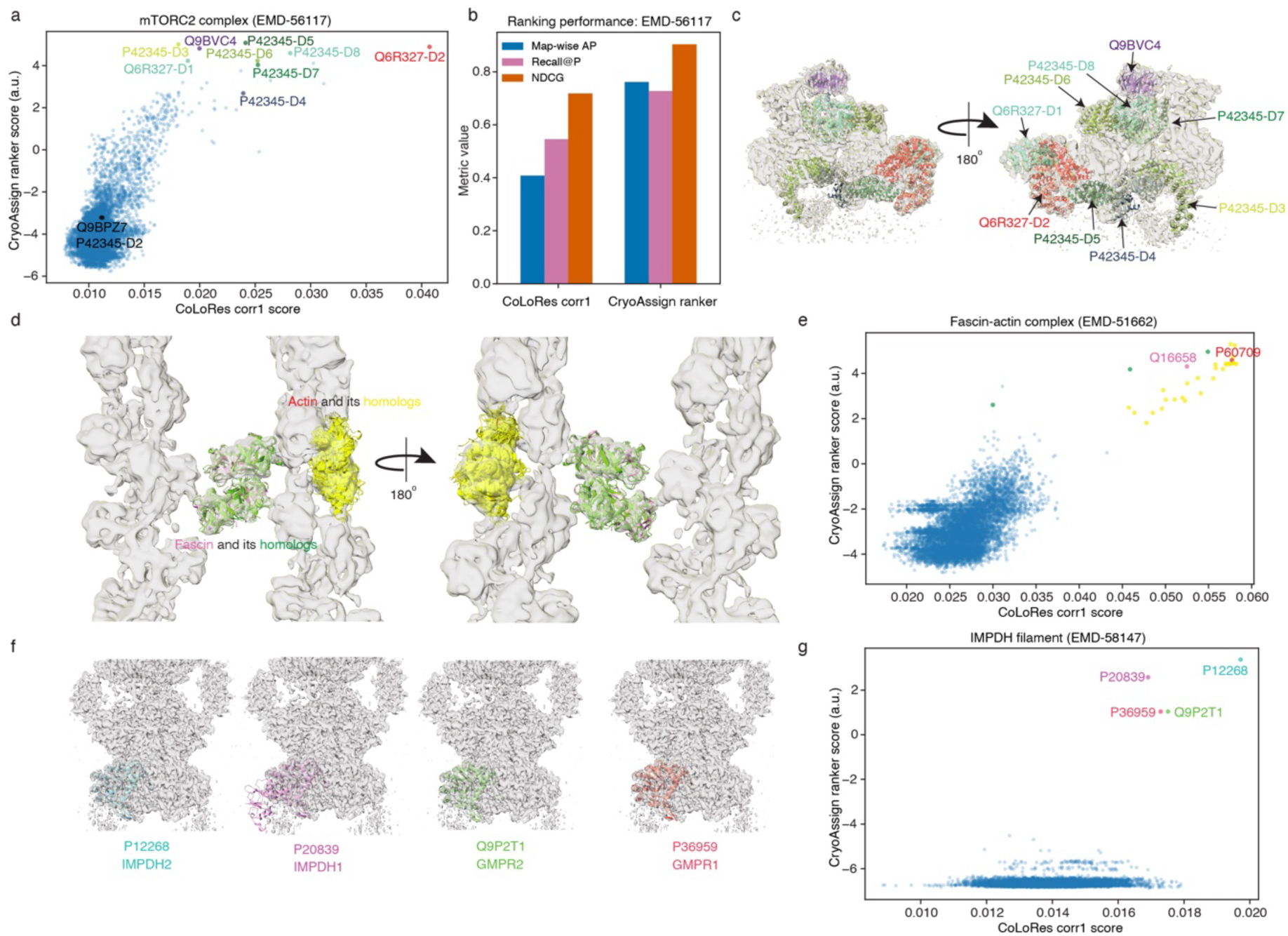
CryoAssign prioritizes candidate proteins from cryo-electron tomography subtomogram-averaged maps. **a,** Scatter plot of CoLoRes corr1 score versus CryoAssign ranker score for the membrane-associated mTORC2 complex (EMD-56117). Correctly identified protein domains are highlighted. **b,** Ranking performance for EMD-56117 comparing the CoLoRes corr1 score and CryoAssign ranker using Map-wise AP, Recall@P, and NDCG. Blue, purple, and vermilion denote Map-wise AP, Recall@P, and NDCG, respectively. **c,** Cryo-ET subtomogram-averaged density map of EMD-56117 overlaid with the correctly identified protein domains. **d,** Cryo-ET subtomogram-averaged density map of the fascin–actin complex (EMD-51662) overlaid with the highest-ranked candidate structures. Actin and its homologs are shown in red and yellow, whereas fascin and its homologs are shown in pink and green. This example illustrates the ambiguity associated with distinguishing proteins with highly similar structural folds at the effective docking resolution. **e,** Scatter plot of CoLoRes corr1 score versus CryoAssign ranker score for EMD-51662. Candidate proteins are color-coded as in **d**. **f,** Cryo-ET subtomogram-averaged density map of filamentous IMPDH from mycophenolic acid-treated HeLa cells (EMD-58147), overlaid separately with the highest-ranked candidate structures. The top-ranked candidates include IMPDH2, IMPDH1, GMPR2, and GMPR1, which adopt closely related structural folds. **g,** Scatter plot of CoLoRes corr1 score versus CryoAssign ranker score for EMD-58147. Candidate proteins are color-coded as in panel **f**. CryoAssign ranker scores in **a**, **e**, and **g** are shown in arbitrary units.

Proteins with highly similar folds remain intrinsically difficult to distinguish by rigid-body docking. Because CryoAssign performs the initial CoLoRes search at the default resolution of 15 Å, docking is driven mainly by overall shape rather than side-chain features, and related architectures can rank together. For EMD-51662^35^ (PDB 9GXI), the top 35 candidates consisted almost exclusively of actin, actin-like proteins, fascin and fascin homologs (**Fig. 5d,e**). This result reflects the intrinsic ambiguity of distinguishing highly similar folds at the effective docking resolution.

A similar ambiguity was observed for an STA map of filamentous IMPDH from mycophenolic acid-treated HeLa cells (EMD-58147^36^). The highest-ranked candidates included IMPDH2, IMPDH1, GMPR2 and GMPR1, all of which have closely related folds (**Fig. 5f,g**). CryoAssign prioritized the correct structural family, but biochemical or contextual information is required to distinguish among these candidates.

### Resolution-dependent performance

To evaluate the effect of map resolution on CryoAssign performance, we selected four high-quality cryo-EM maps and generated low-pass-filtered versions with nominal cutoffs ranging from 4 to 20 Å. Using multiple filtered versions of each parent map enabled a controlled assessment of resolution-dependent performance while holding map identity, candidate database, and search parameters constant. The native and low-pass-filtered maps were searched against the same *E. coli* AlphaFold2-derived candidate database, and performance was evaluated from the ranks of manually defined true-positive candidates.

Across the four datasets, CryoAssign recovered most true hits at high ranks for maps filtered to cutoffs between 4 and 12 Å, although sensitivity to filtering varied among maps (**Fig. 6a–d**). For EMD-9126^37^, all three true hits remained highly ranked from the native resolution of 4.3 Å through a low-pass cutoff of approximately 10 Å (**Fig. 6a**). At approximately 12 Å, one true hit fell to around 50^th^; at 15 Å, a second fell to around 20^th^; and at 20 Å, all three were ranked outside the top 50. For EMD-50723^38^, the single true hit remained highly ranked through 12 Å but declined sharply at 15 Å (**Fig. 6c**). In contrast, performance for EMD-65039^39^ was highly robust, with the true hit remaining at or near the top of the ranked list across all tested filtering conditions (**Fig. 6d**).

**Fig. 6.**
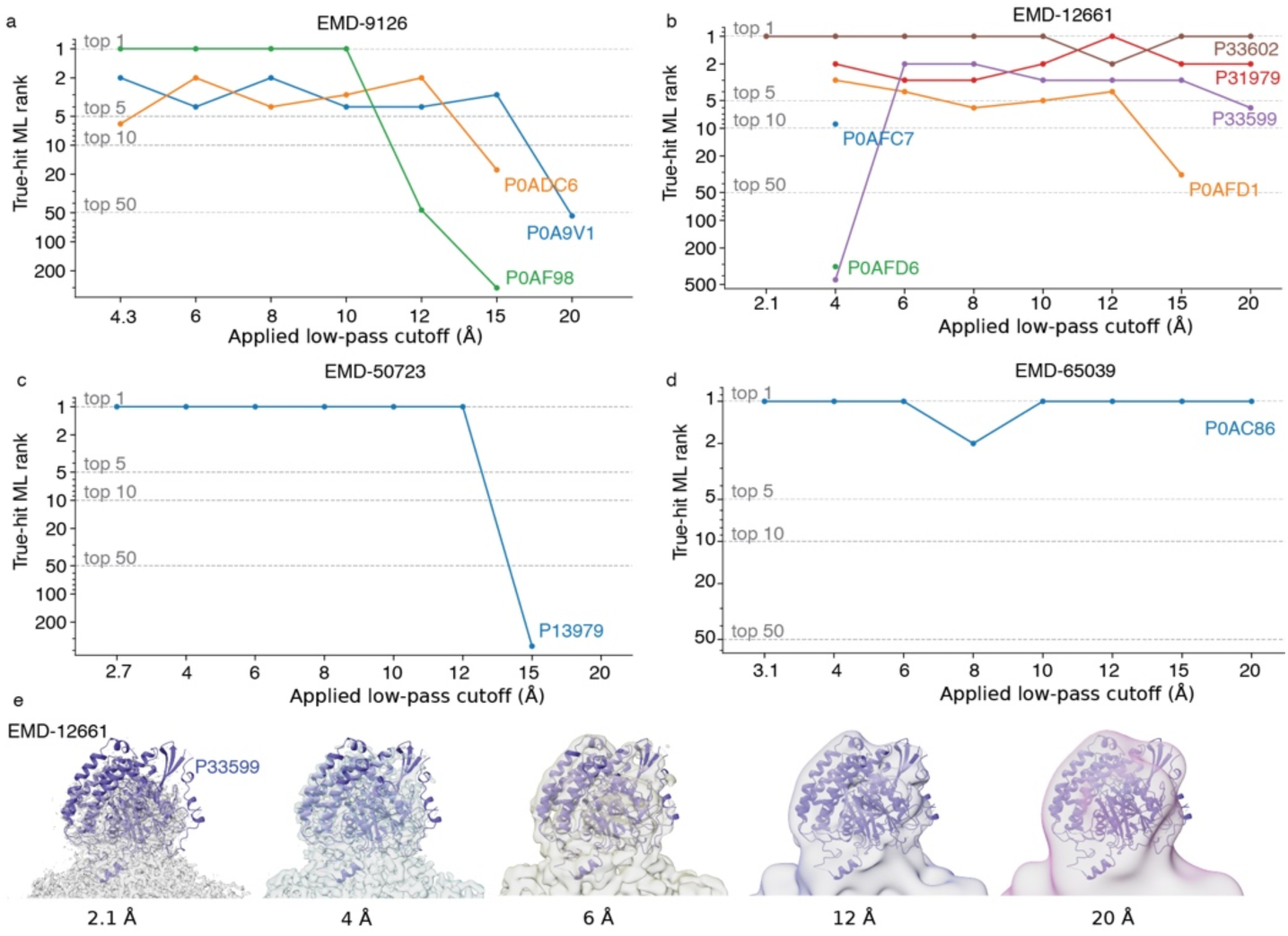
Resolution-dependent performance of CryoAssign under controlled low-pass filtering. **a–d,** True-hit rank trajectories for EMD-9126, EMD-12661, EMD-50723, and EMD-65039, respectively, analyzed in their native forms and after progressive low-pass filtering. The leftmost point in each panel represents the unfiltered native map and is labeled with its reported resolution: 4.3 Å for EMD-9126, 2.1 Å for EMD-12661, 2.7 Å for EMD-50723, and 3.1 Å for EMD-65039. Subsequent points represent the indicated nominal low-pass cutoffs from 4 to 20 Å. For each known true-hit protein, the intact Phenix-processed model and corresponding split-domain candidates were grouped by protein accession, and the candidate with the highest CryoAssign ranker score at each filtering condition was retained. Each colored trajectory therefore represents one protein, although the intact model or split domain contributing the highest score may vary among filtering conditions. The y-axis shows the CryoAssign rank among unique protein entries on an inverted logarithmic scale, with rank 1 at the top. Horizontal reference lines indicate ranks 1, 5, 10, and 50. True-hit proteins are P0AF98, P0ADC6, and P0A9V1 for EMD-9126; P33602, P31979, P33599, P0AFD1, P0AEC7, and P0AFD6 for EMD-12661; P13979 for EMD-50723; and P0AC86 for EMD-65039. Gaps indicate filtering conditions for which no ranked entry was available for the corresponding true-hit protein. The same candidate database and analysis settings were used throughout each filtering series. **e**, EMD-12661 density maps at the indicated native resolution and nominal low-pass cutoffs, overlaid with the P33599 model.

EMD-12661^40^ exhibited a different pattern. In the native 2.1-Å map, only one of the six true hits was ranked within the top 500 candidates (**Fig. 6b**). After low-pass filtering to 4 Å, all six true hits were recovered within the top 500, including four within the top ten. This recovery pattern remained largely stable through a cutoff of approximately 12 Å. At 15 Å, one true hit fell to around 50^th^, although three true hits remained within the top ten even at 20 Å. The most pronounced improvement was observed for P33599, which rose from approximately 500^th^ at the 4-Å cutoff to 2^nd^ at 6 Å and remained within the top ten through 20 Å. Inspection of the corresponding density suggested that the region containing P33599 was oversharpened and fragmented in the native map (**Fig. 6e**). Progressive low-pass filtering produced more continuous density features, which likely contributed to the improved ranking.

Together, these results indicate that CryoAssign performed most consistently at applied low-pass cutoffs of approximately 4–12 Å, whereas rankings generally became less reliable beyond approximately 15 Å. The separation between scores of true-positive and negative candidates narrowed at coarser cutoffs (**Supplementary Fig. 2**). However, the contrasting behavior of EMD-65039 and EMD-12661 shows that nominal resolution alone does not determine performance. Local map quality, density continuity, sharpening and preservation of overall shape also influence recovery.

## Discussion

In this study, we developed CryoAssign, an automated pipeline that combines CoLoRes rigid-body docking with a learning-to-rank framework for identifying unknown proteins from cryo-EM density maps. By integrating complementary docking-derived features, CryoAssign improved candidate prioritization relative to the original CoLoRes unnormalized cross-correlation score and enabled assignment of previously unexplained density maps from *E. coli*, human extracellular vesicle, and cryo-ET datasets.

A key aspect of CryoAssign is the formulation of protein identification as a learning-to-rank problem. This more naturally reflects the objective of prioritizing correct candidates from large structural databases than binary classification. It also accommodates maps with multiple true components, for which all correct candidates should be ranked highly. Accordingly, we evaluated performance using Recall@P, map-wise AP and NDCG, which quantify how effectively true hits are concentrated near the top of the ranked candidate list.

The engineered features describe distinct aspects of CoLoRes docking, including score separation, regional real-space correlation, overlap coherence, structural continuity and candidate size. Individual features improved candidate prioritization for subsets of benchmark maps, but their relative performance varied. Integrating these descriptors produced more robust overall performance than any individual feature, with the trained model achieving the strongest or near-strongest per-map performance across map-wise AP, Recall@P and NDCG. Further improvements are likely as larger and more diverse training datasets become available.

CryoAssign also differs from related approaches in its input requirements and operating regime. Unlike sequence-based identification approaches such as CryoID^41^ or model-building approaches such as ModelAngelo, CryoAssign identifies proteins through large-scale structure-based matching and does not require near-atomic side-chain features. Because CoLoRes uses a default target resolution of 15 Å, the initial rigid-body search is driven primarily by overall molecular shape and map–model agreement rather than side-chain density. This makes CryoAssign applicable to intermediate-resolution cryo-EM maps and cryo-ET subtomogram averages, where side-chain features are often unresolved or unreliable. In addition, unlike DomainFit, which relies on segmented density regions, CryoAssign performs exhaustive searches against intact density maps without prior segmentation. This avoids the need to define component boundaries before searching and makes the workflow broadly applicable to cases in which the number, boundaries, or identities of molecular components are unknown.

A current limitation of CryoAssign is that the learning-to-rank model relies on features derived from CoLoRes docking and therefore depends on the quality of the initial rigid-body placement. Although the exhaustive six-dimensional search reduces the risk of missing the correct solution, docking can still be affected by map quality, conformational variability, structural symmetry and limited shape information. Consequently, correct candidates may receive unexpectedly low rankings when their placements are suboptimal. In EMD-22475 and EMD-56117, four and two correct candidates, respectively, ranked well below their expected positions because CoLoRes did not place them optimally (**Figs. 4a and 5a**). These examples illustrate that current performance is partly limited by the initial docking solution. CoLoRes can be run at finer target resolutions and with smaller search step sizes, but at increased computational cost. Future improvements could incorporate features from multiple docking algorithms or flexible-fitting approaches, reducing dependence on a single rigid-body docking solution.

Overall, CryoAssign reframes large-scale identification of unknown proteins from cryo-EM and cryo-ET density maps as a learning-to-rank problem and translates this framework into an automated, scalable pipeline. By integrating structure-database preparation, exhaustive docking, feature extraction and data-driven prioritization, CryoAssign enables proteome-scale searches of intact, intermediate-resolution maps without prior segmentation. Expansion of the training data and advances in map reconstruction, structure prediction, docking and machine learning should further improve its accuracy and applicability.

## Methods

### Structure database preparation

To support CryoAssign searches, we developed a parallelized download script, download_structures_parallel2.py, to retrieve coordinate files from either the AlphaFold Protein Structure Database (AFDB) or the Research Collaboratory for Structural Bioinformatics (RCSB) PDB. Species names or proteome identifiers were used to obtain complete structure sets for a given proteome. For AlphaFold models, we downloaded 23,586 models for *Homo sapiens* (UP000005640) and 4,370 models for *Escherichia coli* (UP000000625) from AFDB.

### Domain splitting of AlphaFold2-predicted models

Because the relative orientations of domains in full-length AlphaFold2-predicted models may not reflect the conformations captured in experimental density maps, the models were split into individual domains before docking. Domain splitting was performed using phenix.process_predicted_model^17^ from the Phenix package^18^ through a wrapper script, process_predicted.py, which passes PAE files and associated metadata when available. For all downloaded AlphaFold2 models, we used a minimum pLDDT cutoff of 50, a PAE cutoff of 10 Å, and a PAE power parameter of 1.

### Domain and PDB fitting using CoLoRes

For CryoAssign screening, each candidate structure was docked independently into the target cryo-EM map using CoLoRes^14^ from the Situs package^13^. The alignment step was executed in batch mode, with one CoLoRes run performed per candidate model and the resulting poses written to candidate-specific output directories. CoLoRes was run using Laplacian cross-correlation scoring (-corr 1), a target resolution of 15 Å (-res 15.0), and a density cutoff of 0.0 (- cutoff 0.0). CPU allocation was controlled at the batch level, with the number of processes per docking run (-nprocs) matched to the computational resources assigned by the HPC scheduler. For each candidate, the resulting col_*.pdb outputs were collected into per-candidate folders and used for downstream feature extraction, reranking, and selection of top-scoring fits.

### Computational performance

The computational cost of CryoAssign depends primarily on the size of the target density map, the search resolution, and the number of candidate structures. As a representative example, searching the human AlphaFold2 database (40,596 processed domains) against an approximately 1-MDa density map using the default CoLoRes search resolution of 15 Å required approximately 48 hours on 288 CPU cores (72 concurrent tasks with 4 cores per task).

### Docking-derived features

For each candidate structure, CoLoRes output files were parsed to derive a compact set of features describing five complementary aspects of docking quality: (i) docking-score separation, (ii) map–model agreement, (iii) density coverage, (iv) structural continuity, and (v) candidate size (**Supplementary Table 1**).

The primary CoLoRes score (corr1) and the score-separation feature gap12 quantify the strength and uniqueness of the best-ranked docking solution. Map-model agreement was characterized using three masked Pearson correlation coefficients calculated over different spatial regions of the density map, including the strict overlap region, a locally expanded overlap region, and the complete model mask. All correlation-based features were calculated using the masked Pearson correlation coefficient,

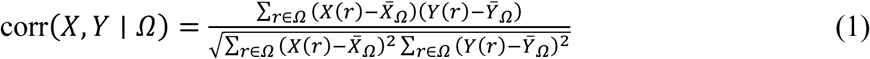

where X(r) and Y(r) denote the experimental and model-derived densities, respectively, at voxel r; Ω is the voxel set over which the comparison is performed; and X̅_Ω_ and Ȳ_Ω_ are the corresponding mean densities within Ω.

Additional features were designed to quantify the extent and spatial coherence of the docking solution, including the size of the largest connected component relative to the model mask, the degree of overlap fragmentation, the number of significant map voxels explained by the candidate, the fraction of residues supported by experimental density, and the total number of atoms in the candidate structure. Together, these features summarize complementary properties of the docking solution that are subsequently integrated by the learning-to-rank model.

### Training learning-to-rank models

Because the practical objective of CryoAssign is to prioritize correct candidates near the top of a ranked list rather than classify candidates individually as correct or incorrect, candidate prioritization was formulated as a learning-to-rank problem.

Docking-derived feature tables were generated for 122 training maps and 22 held-out benchmark maps. Each map was docked against the corresponding species-specific candidate structure database generated from AlphaFold2-predicted structures, and docking-derived features were extracted for every candidate. The cryo-EM maps were partitioned at the map level before model training, ensuring that all docking results derived from a given map were assigned exclusively to either the training or held-out benchmark set and preventing information leakage during model fitting. Among these maps, 108 training maps and 14 held-out benchmark maps contained at least one manually confirmed true-positive candidate and were used for recovery-oriented metric calculations.

Ground-truth labels were assigned by manual inspection of the docking results using the corresponding experimentally determined structures deposited in the PDB as references. Candidate docking solutions were labeled as positive examples only when they represented the correct protein or protein domain and the top-ranked CoLoRes pose correctly reproduced the experimentally observed placement within the density map. All remaining candidates were labeled as negative examples.

Supervised learning-to-rank models were trained using XGBRanker version 3.2.0 with the rank:ndcg objective. Candidate lists were grouped by map so that each cryo-EM map represented one ranking group. No additional hyperparameter optimization was performed; default XGBoost hyperparameters were used apart from the specified ranking objective. Similar performance was observed across several predefined feature sets, indicating that the improvement was not dependent on a single narrowly optimized feature combination. The orig_core_mixed feature set was selected for the final model because it retained interpretable features describing docking-score separation, map–model agreement, structural continuity, density coverage, candidate size, and the original CoLoRes score. This final feature set comprised ten features: corr1, gap12, overlap_cc_strict_orig, overlap_cc_local_orig, masked_realspace_cc_orig, supported_residue_fraction_z, largest_overlap_component_fraction_model, overlap_fragmentation_penalty, map_explained_voxels, and n_atoms_total.

Model performance was evaluated on the 14 held-out benchmark maps containing at least one manually confirmed true-positive candidate using three primary ranking metrics: Recall@P, map-wise average precision (AP) and NDCG. Secondary summary metrics, including MRR,

Top-1, Top-5 and Top-10 hit rates and mean positive rank, are reported in **Supplementary Table 2**. For visual evaluation of the training results, **Supplementary Figure 1** was generated programmatically in Python using Matplotlib^42^.

For a map containing P positive candidates, Recall@P was defined as

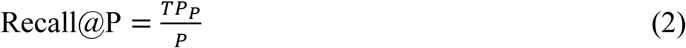

where TP_P_ is the number of true-positive candidates recovered within the top P ranked candidates.

Average precision (AP) for an individual map was calculated as

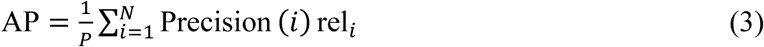

where N is the total number of ranked candidates and rel_i equals 1 for a true-positive candidate and 0 otherwise. Precision(i) is the precision at rank i,

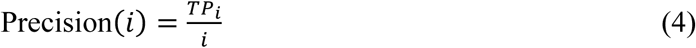

TP_i is the number of true-positive candidates within the top i ranked candidates. The reported map-wise AP was calculated as the mean AP across the 14 held-out maps used for recovery-oriented evaluation.

The NDCG was computed using binary relevance labels. The discounted cumulative gain (DCG) was calculated as

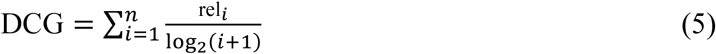

where i is the rank position and rel_i equals 1 for a true-positive candidate and 0 otherwise. NDCG was then calculated as

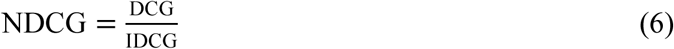

where IDCG denotes the ideal discounted cumulative gain obtained from the optimal ranking of the same set of candidates, in which all true-positive candidates precede all false-positive candidates,

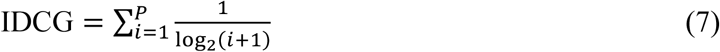

Secondary metrics reported in **Supplementary Table 2** were mean reciprocal rank (MRR), Top-1, Top-5 and Top-10 hit rates, and mean positive rank. MRR was the mean reciprocal rank of the first positive candidate across maps. The Top-k hit rate was the fraction of maps with at least one positive candidate within the top k ranks. Mean positive rank was calculated within each map and then averaged across maps.

### CryoAssign analysis of unknown *E. coli* protein maps

Two unknown cryo-EM density maps were obtained from a recombinant protein sample expressed in *Escherichia coli*. Candidate structures were generated by downloading the complete *E. coli* AlphaFold2 proteome (UniProt proteome UP000000625) and processing the predicted models using phenix.process_predicted_model, yielding 11,750 candidates. Each density map was searched independently against this database using CryoAssign, and candidate structures were prioritized using the trained learning-to-rank model. All large-scale searches were performed on a high-performance computing (HPC) cluster.

### CryoAssign analysis of unknown human protein map

An unknown cryo-EM density map obtained from human extracellular vesicle samples was searched against a human AlphaFold2-derived candidate structure database. The database was generated by processing the complete human AlphaFold2 proteome (UniProt proteome UP000005640) using phenix.process_predicted_model, yielding 40,596 candidate domains. Candidate structures were docked using CoLoRes, and the resulting docking solutions were prioritized using the trained learning-to-rank model.

### CryoAssign analysis of the radial spoke complex from the *Chlamydomonas reinhardtii* axoneme

A total of 16,153 AlphaFold2-predicted protein structures for *Chlamydomonas reinhardtii* were downloaded and processed using phenix.process_predicted_model. The resulting candidate structure database comprised 35,479 entries, including both split domains and intact protein models. The radial spoke complex cryo-EM map (EMD-22475) was searched against this database using CryoAssign, and candidate structures were prioritized using the trained learning-to-rank model. The resulting rankings were compared with the experimentally assigned protein components reported for the corresponding structure.

### CryoAssign analysis of cryo-ET subtomogram-averaged maps

Cryo-electron tomography subtomogram-averaged maps, including EMD-51662^35^, EMD-56117^34^ and EMD-58147^36^, were downloaded from the Electron Microscopy Data Bank (EMDB)^43^. Each map was searched against the corresponding species-specific AlphaFold2-derived candidate structure database generated using phenix.process_predicted_model. Candidate structures were docked using CoLoRes, and the resulting docking solutions were prioritized using the trained learning-to-rank model. These datasets were used to evaluate the applicability of CryoAssign to cryo-ET maps spanning a range of resolutions and structural complexities.

### Resolution-dependent performance of CryoAssign

Cryo-EM maps EMD-9126^37^, EMD-50723^38^, EMD-65039^39^, and EMD-12661^40^ were analyzed in their native forms and after low-pass filtering to nominal cutoffs of 4, 6, 8, 10, 12, 15, and 20 Å using the RELION utility relion_image_handler^2^. The 4-Å filtering condition was omitted for EMD-9126 because its reported native resolution was 4.3 Å. For EMD-50723 and EMD-12661, masked maps generated from the corresponding half-maps were used to reduce contributions from background noise before low-pass filtering.

Each native or low-pass-filtered map was searched against the same candidate database comprising 11,750 Phenix-processed entries derived from the complete *E. coli* AlphaFold2 proteome (UniProt proteome UP000000625). The same CoLoRes search parameters, feature-extraction settings, and trained learning-to-rank model were used for all filtering conditions. For EMD-50723, the AlphaFold2 model of the known true-hit protein P13979 was additionally downloaded, processed using the same procedure, and added to the candidate database. The resulting docking solutions were prioritized using the fixed trained learning-to-rank model.

### Cryo-EM data collection and processing

Cryo-EM data for DnaK and ArnA were collected on a Thermo Fisher Scientific Titan Krios G2 electron microscope operated at 300 kV and equipped with a Gatan K3 direct electron detector and a Gatan Imaging Filter with a 20-eV energy-selecting slit. Movies were recorded in super-resolution mode at a nominal magnification of 130,000×, corresponding to a pixel size of 0.664 Å per pixel. Each movie was acquired with a total electron exposure of 50 e^-^ Å^-2^, fractionated over 50 frames. Defocus values ranged from −0.5 to −3.5 μm.

Data were processed using cryoSPARC v4.1.1^3^. Two datasets comprising 12,002 and 10,092 movies, respectively, were subjected independently to patch motion correction and estimation of contrast transfer function parameters. For dataset 1, initial blob picking from 200 micrographs yielded 92,100 particles. Two-dimensional class averages generated from these particles were used as templates for particle picking across the complete dataset. Following two rounds of 2D classification, 362,022 large-particle images and 23,090 small-particle images were retained. For dataset 2, initial blob picking from 200 micrographs yielded 94,533 particles, which were used to generate templates for particle picking across the complete dataset. Following two rounds of 2D classification, 390,704 large-particle images and 110,178 small-particle images were retained.

The retained particle images from the two datasets were combined, yielding stacks of 752,726 large-particle images and 132,536 small-particle images for subsequent 3D reconstruction.

For the small particles corresponding to DnaK, one round of heterogeneous refinement retained 41,666 particles. These particles were subjected to non-uniform refinement followed by local refinement, yielding a final reconstruction at 4.2 Å resolution.

For the large particles corresponding to ArnA, initial 3D classification and refinement yielded reconstructions in which only one or two blades of the three-bladed propeller were well resolved. To improve the reconstruction, the combined particle stack was exported to RELION 4^2^ for alignment-free masked 3D classification. Two masks covering the poorly resolved regions were used. This classification identified a subset of 150,866 particles with well-resolved density for all three blades. Subsequent non-uniform and local refinement with C3 symmetry yielded a final reconstruction at 3.9 Å resolution. C3 symmetry was imposed during refinement to avoid enforcing the additional twofold symmetry because of conformational heterogeneity.

Cryo-EM data for α2-macroglobulin were collected on a Thermo Fisher Scientific Glacios 2 electron microscope operated at 200 kV and equipped with a Falcon 4i direct electron detector and a Selectris X energy filter with a 10-eV slit. Movies were recorded at a nominal magnification of 130,000×, corresponding to a pixel size of 0.88 Å per pixel. Each movie was acquired with a total electron exposure of 42 e^-^ Å^-2^, fractionated over 30 frames. Defocus values ranged from −0.4 to −2.0 μm.

Data were processed using cryoSPARC v4.7.1. A dataset comprising 11,303 movies was binned twofold and subjected to patch motion correction and estimation of contrast transfer function parameters, resulting in micrographs with a pixel size of 1.76 Å. Initial blob picking from 200 micrographs yielded 14,103 particles. Two-dimensional class averages generated from these particles were used as templates for particle picking across the complete dataset. Following two rounds of 2D classification, 249,772 particles were retained for 3D reconstruction. One round of heterogeneous refinement retained 68,118 particles, which were subjected to non-uniform refinement with D2 symmetry, yielding a final reconstruction at 4.5 Å resolution.

For all structures, the reported resolution was estimated using the gold-standard Fourier shell correlation criterion of 0.143. Cryo-EM data collection and refinement statistics are provided in **Supplementary Table 3**.

### Use of generative AI

Generative AI (ChatGPT Enterprise, OpenAI) was used during the preparation of this work to assist with code development, interpretation of computational outputs, and manuscript editing.

## Data availability

Cryo-EM density maps of DnaK, ArnA, and α2-macroglobulin have been deposited in the Electron Microscopy Data Bank under accession codes EMD-78960, EMD-78961, and EMD-78963, respectively.

## Code availability

The CryoAssign code is available via Zenodo at https://zenodo.org/records/22434065?preview-1=token-eyJhbGciOiJIUzUxMiJ9.eyJpZCI6ImVjMjhlNmU2LWEzNTAtNGYwNy04NGE3LTkwY2FhMjRhZTJkNSIsImRhdGEiOnt9LCJyYW5kb20iOiJkM2IxZjEyMzFiMTczYzE2NTFhNzdmYzFlMGViNjY4ZiJ9.gO2eomLq36mh4zkxoHmcmklK_nL3gACQMLtPNZDi8gmLXjzSFMDTXXNazGeoqPWEIkmslEB_SgA71VXyBQIH7w

A tutorial data set is available via Zenodo at https://zenodo.org/records/22397582?preview-1=token-eyJhbGciOiJIUzUxMiJ9.eyJpZCI6IjhmYzg0MTUzLWJkOTktNDJjMi05NTdmLWM3NjEyZWVhYWU5ZSIsImRhdGEiOnt9LCJyYW5kb20iOiJjYTAwNTdmYzFlMGMwOGRjYjFhYjhjYTNiODNkYWY2YyJ9.Z-13qkvLps9W-FDT_Z74yYCaaNJ2cDZr0AISBBirGj9lI12UmY5A44iqzEWCSWMlw0uWmr6BoBblVGgC8hv82g

## Acknowledgements

Cryo-EM data were collected at the Structural Biology Core at Mayo Clinic and at the Hormel Institute. We thank Janarjan Bhandari of the Hormel Institute for assistance with data collection.

## Author contributions

Q.H. and G.M. conceptualized the study. Q.H. conducted the investigation and wrote the original draft. Q.H. and G.M. reviewed and edited the manuscript.

## Additional Information

**Supplementary information** The online version contains supplementary material, including two Supplementary Figures, three Supplementary Tables, and one Supplementary Movie.

