## Supplementary Information for "A learning-to-rank framework for large-scale identification of unknown proteins from cryo-EM and cryo-ET maps"

Qi Hu[1,2] & Georges Mer[1,2]

[1]Department of Biochemistry and Molecular Biology, Mayo Clinic, Rochester, MN, USA

[2]Structural Biology Core, Mayo Clinic, Rochester, MN, USA

#### Supplementary Table 1 | Docking-derived features used in the final CryoAssign learning-to-rank model.

| Feature | Feature type | Description | Formula |
|---|---|---|---|
| corr1 | Docking score | CoLoRes unnormalized cross-correlation score | CoLoRes output |
| gap12 | Score separation | Difference between the top two CoLoRes scores | $s_1 - s_2$ |
| overlap_cc_strict_orig | Map-model agreement | Pearson correlation within the strict overlap region | Eq. (1), Ω = Ω_strict |
| overlap_cc_ local_orig | Map-model agreement | Pearson correlation within the dilated overlap region | Eq. (1), Ω = Ω_local |
| masked_realspace _cc_orig | Map-model agreement | Pearson correlation over the complete model mask | Eq. (1), Ω = Ω_model |
| map_explained _voxels | Density coverage | Number of significant map voxels explained by the candidate | \|Ω_map ∩ Ω_model\| |
| supported_ residue_fraction_z | Density coverage | Fraction of residues supported by experimental density | N_res,supported / N_res |
| largest_overlap_ component_ fraction_model | Structural continuity | Size of the largest connected overlap component normalized by the model-mask size | \|C_max\| / \|Ω_model\| |
| overlap_fragmentation _penalty | Structural continuity | Fragmentation penalty of the overlap region | 1 − \|C_max\| / \|Ω_overlap\| |
| n_atoms_total | Candidate size | Total number of atoms in the candidate structure | N_atoms |

s1 and s2 denote the highest and second-highest CoLoRes scores, respectively. Ω_strict, Ω_local and Ω_model denote the strict overlap region, the dilated overlap region and the model mask, respectively. Ω_map denotes the significant map mask, and Ω_overlap = Ω_map ∩ Ω_model denotes the overlap mask. |Ω| denotes the number of voxels in the corresponding mask. C_max denotes the largest connected component of Ω_overlap. N_res denotes the total number of residues in the candidate structure, and N_res,supported denotes the number of supported residues. A residue is considered supported if at least 50% of its sampled robust density values satisfy z ≥ z0. A threshold of z0 = 1.0 was used. N_atoms denotes the total number of atoms in the candidate structure.

#### Supplementary Table 2 | Performance comparison between the CoLoRes corr1 baseline and the final CryoAssign ranker on the held-out benchmark set.

| | CoLoRes corr1 baseline | CryoAssign ranker |
|---|---|---|
| Map-wise AP | 0.8420 | 0.9813 |
| **MRR** | 0.9464 | 1.0000 |
| **Recall@P** | 0.8095 | 0.9583 |
| **NDCG** | 0.9134 | 0.9940 |
| **Top-1 hit** | 0.9286 | 1.0000 |
| **Top-5 hit** | 1.0000 | 1.0000 |
| **Top-10 hit** | 1.0000 | 1.0000 |
| **Mean positive rank** | 12.70 | 2.93 |

Metrics were calculated for held-out map groups containing at least one positive candidate.

**Recall@P** denotes the fraction of positive candidates recovered within the top *P* positions, where *P* is the number of positive candidates for that map.

**Top-1, Top-5, and Top-10 hit rates** denote the fraction of maps for which at least one positive candidate was ranked within the top 1, 5, or 10 positions, respectively.

**Mean positive rank** is the mean rank of all positive candidates within each map, averaged across maps; lower values indicate better ranking performance.

**MRR**, mean reciprocal rank; $MRR = (1/N)\Sigma(1/r_i)$, where $r_i$ is the rank of the first true candidate for map i.

**NDCG**, normalized discounted cumulative gain; AP, average precision.

**Supplementary Table 3 | Cryo-EM data collection, refinement and validation statistics**

| | DnaK | ArnA | A2M |
|---|---|---|---|
| **Data collection and processing** | | | |
| Magnification | 130,000 | 130,000 | 165,000 |
| Voltage (kV) | 300 | 300 | 200 |
| Electron exposure ($e^{-}$ Å$^{-2}$) | 50 | 50 | 50 |
| Defocus range (μm) | −0.5 to −2.5 | −0.5 to −2.5 | −0.4 to −2.0 |
| Pixel size (Å) | 0.664 | 0.664 | 1.76 |
| Symmetry imposed | C1 | C3 | D2 |
| Initial particle images (no.) | 132,536 | 752,726 | 249,772 |
| Final particle images (no.) | 41,666 | 150,866 | 68,118 |
| Map resolution (Å) | 4.2 | 3.9 | 4.5 |
| FSC threshold | 0.143 | 0.143 | 0.143 |
| **Refinement** | | | |
| Map sharpening B factor (Å$^2$) | −145 | −120 | −160 |

**Supplementary Figure 1 | Per-map performance of CryoAssign on the held-out benchmark set.** Each page summarizes the performance of CryoAssign for one of the 14 held-out benchmark maps containing at least one manually confirmed true-positive candidate. For each map, score distributions, feature relationships, cumulative recovery curves, and per-map ranking metrics are shown. Positive candidates correspond to manually annotated true hits.

Supplementary Figure 1a

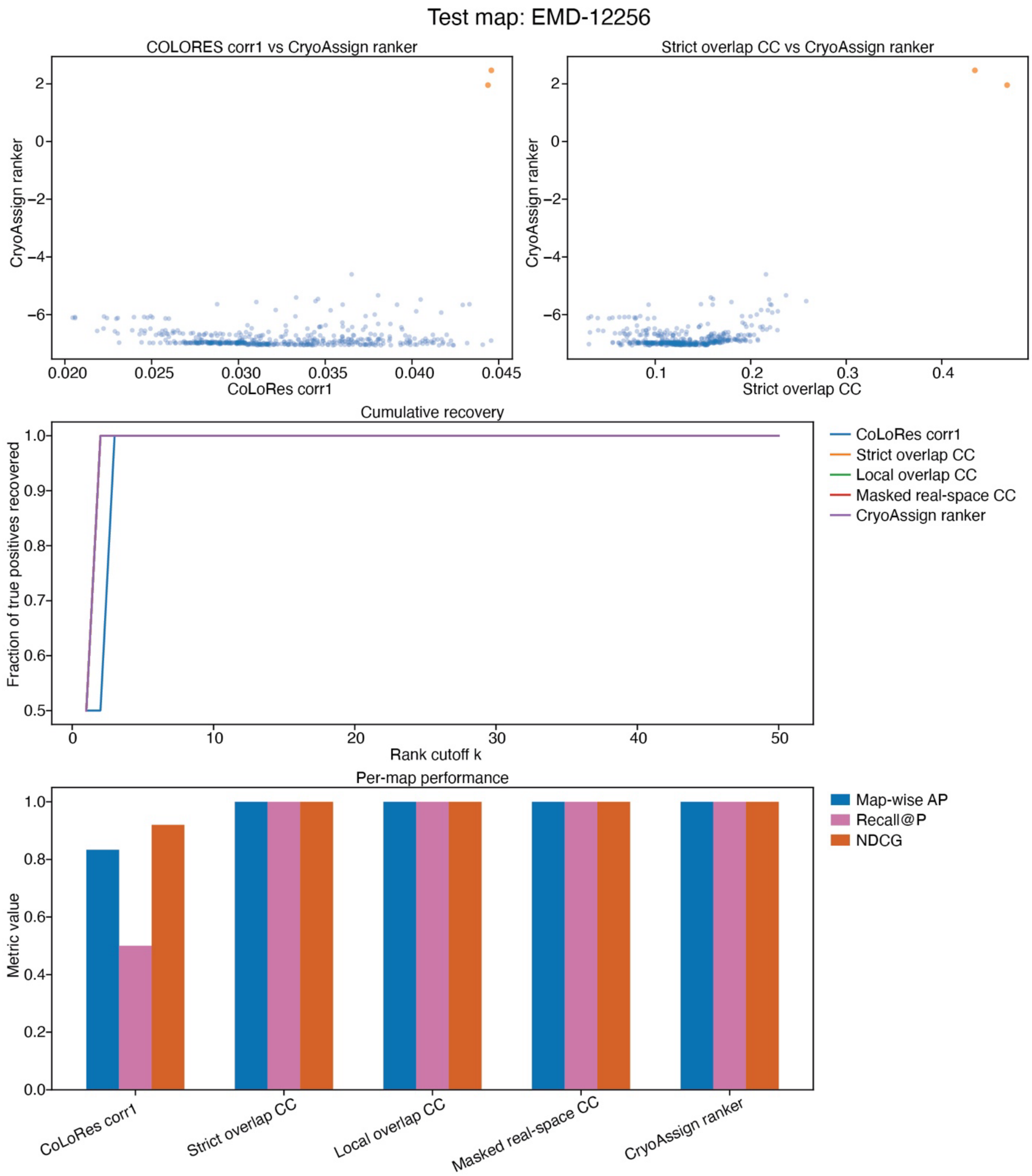

#### Supplementary Figure 1b

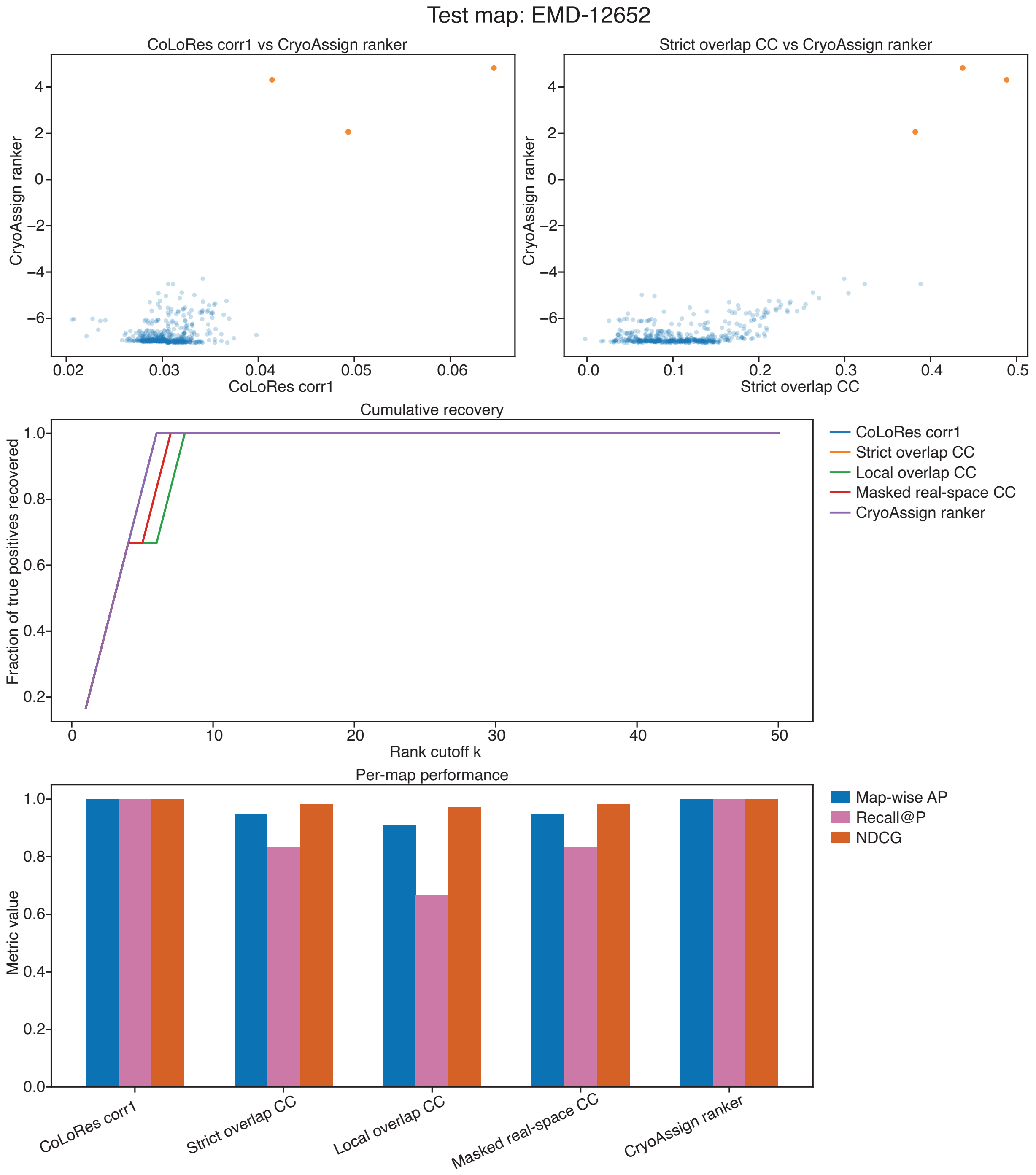

### Supplementary Figure 1c

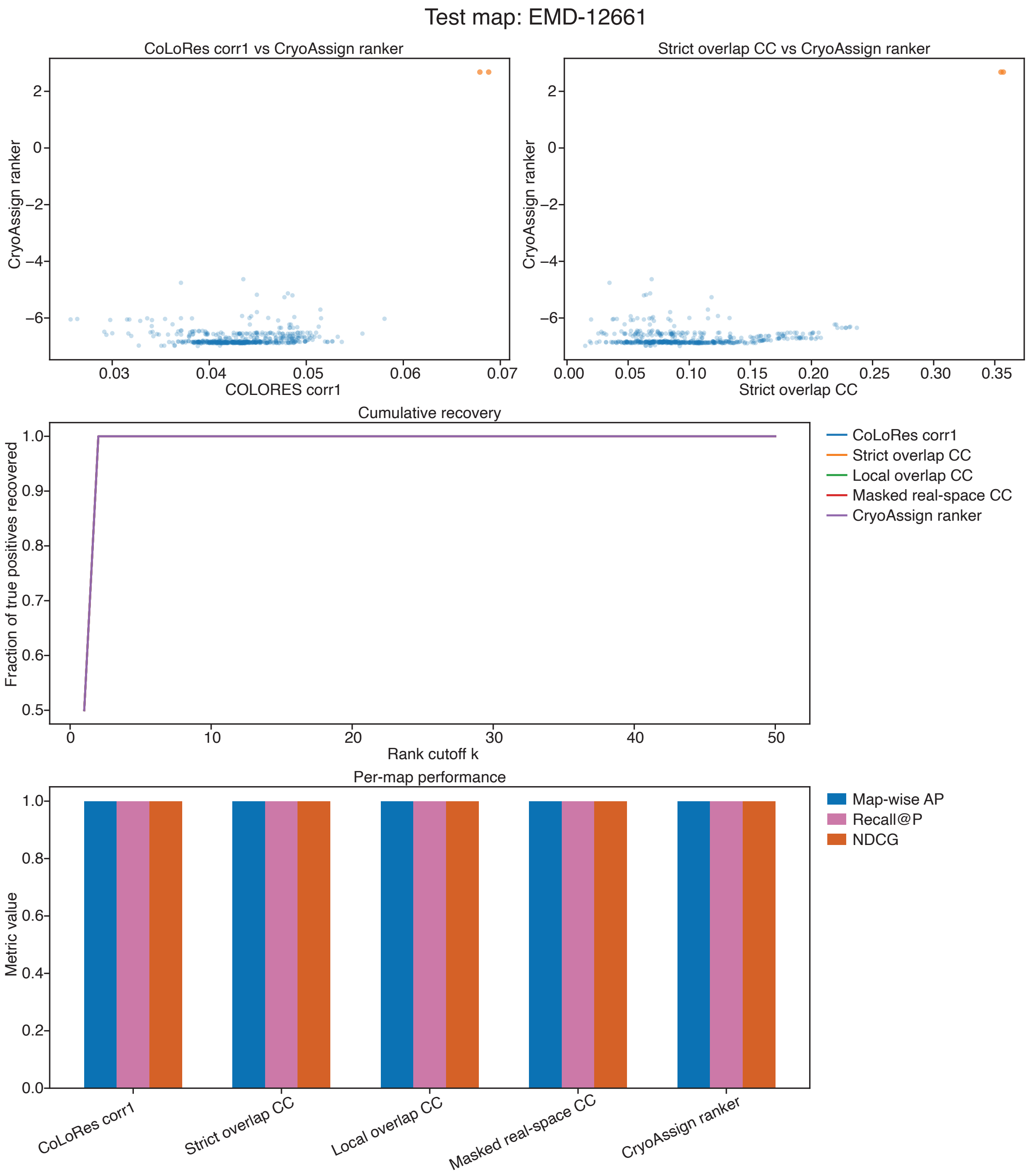

### Supplementary Figure 1d

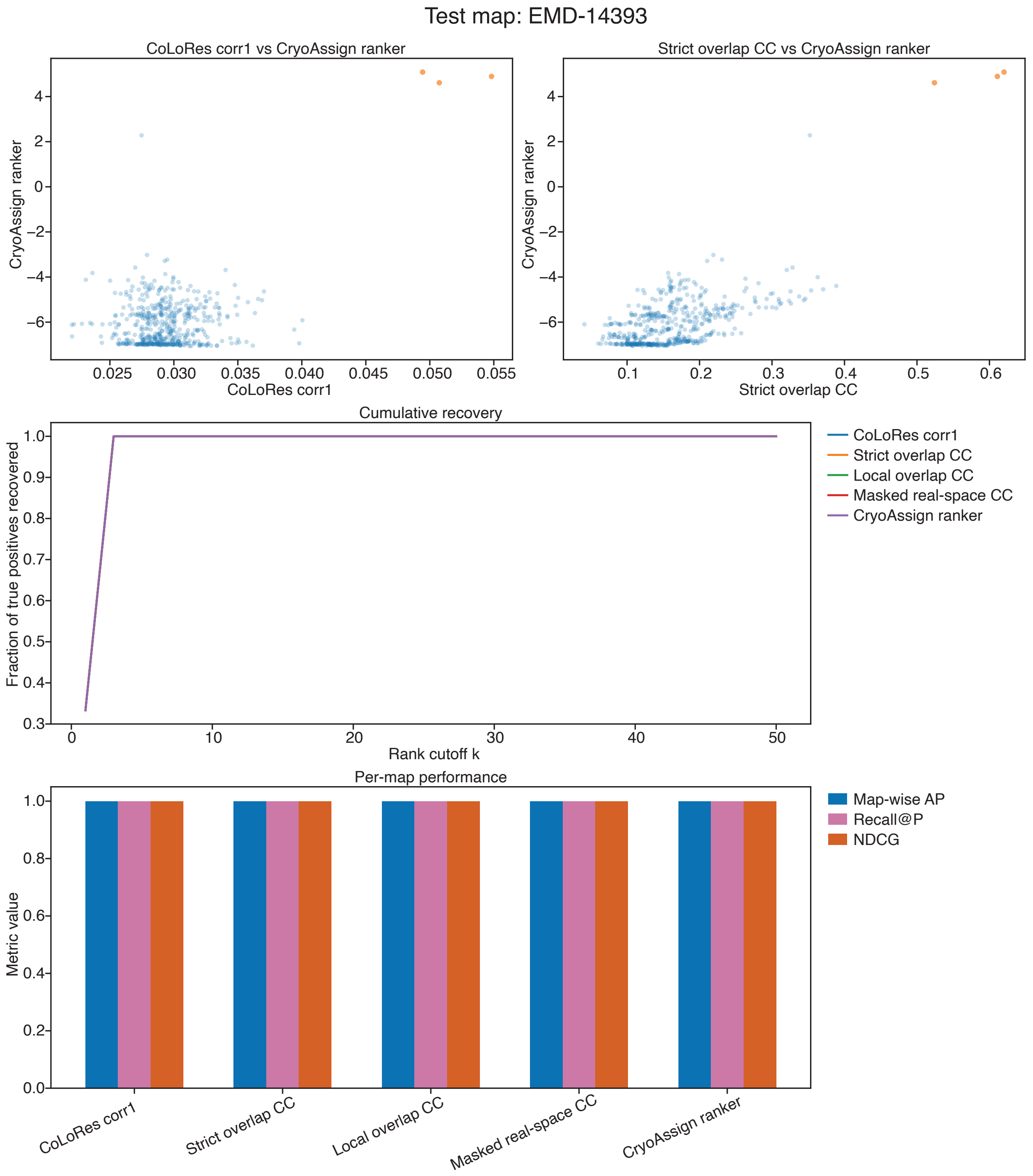

### Supplementary Figure 1e

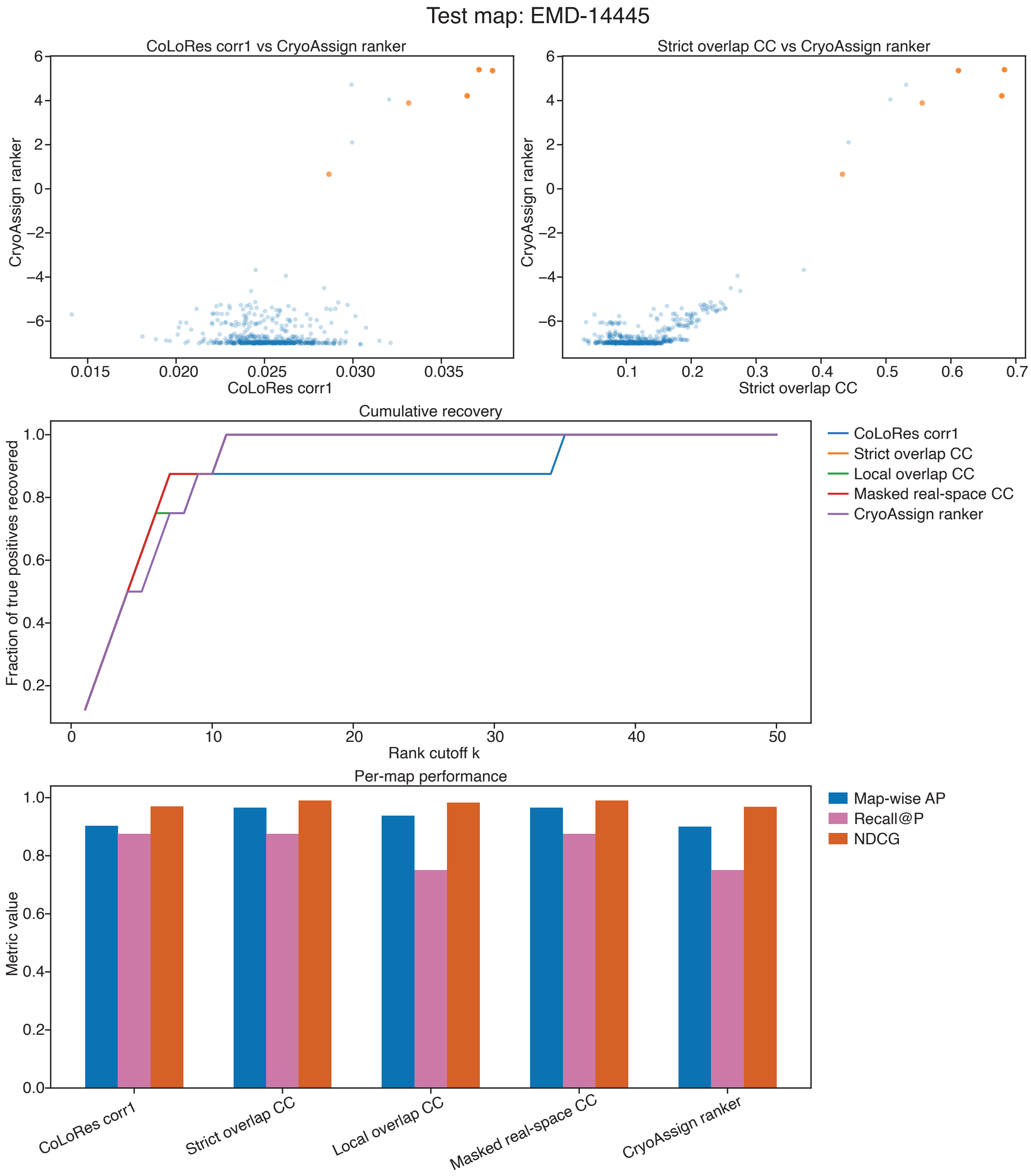

### Supplementary Figure 1f

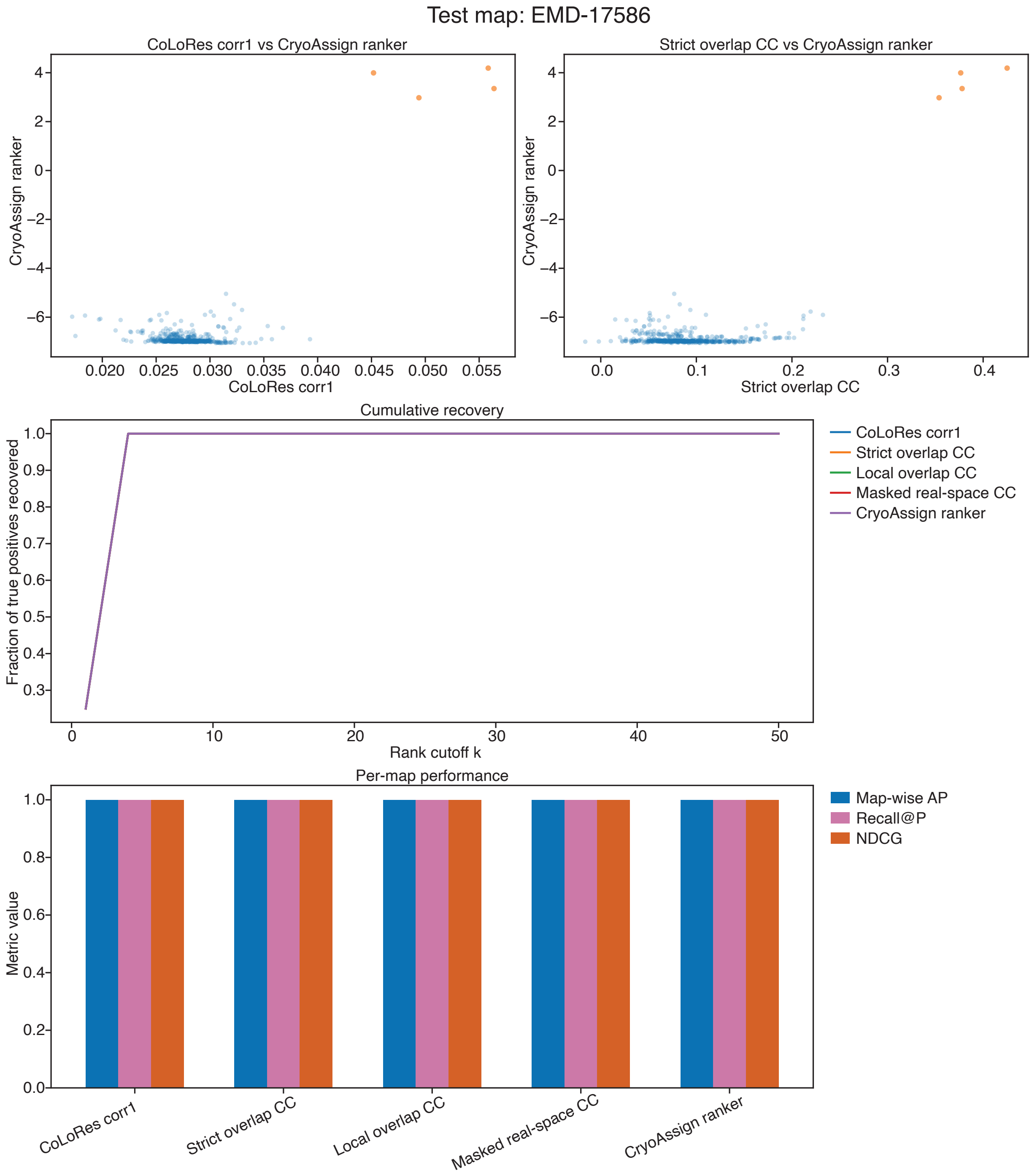

### Supplementary Figure 1g

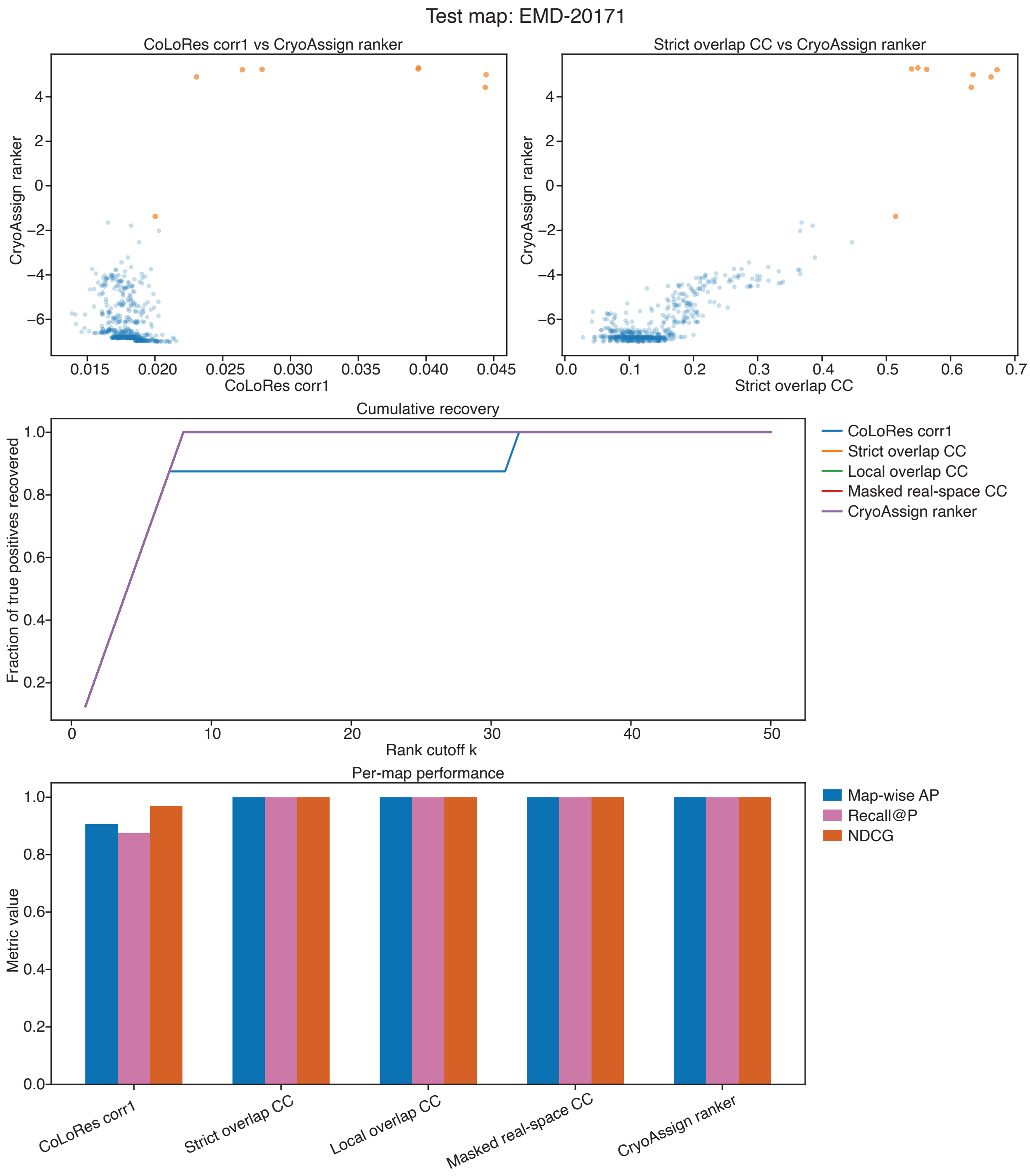

### Supplementary Figure 1h

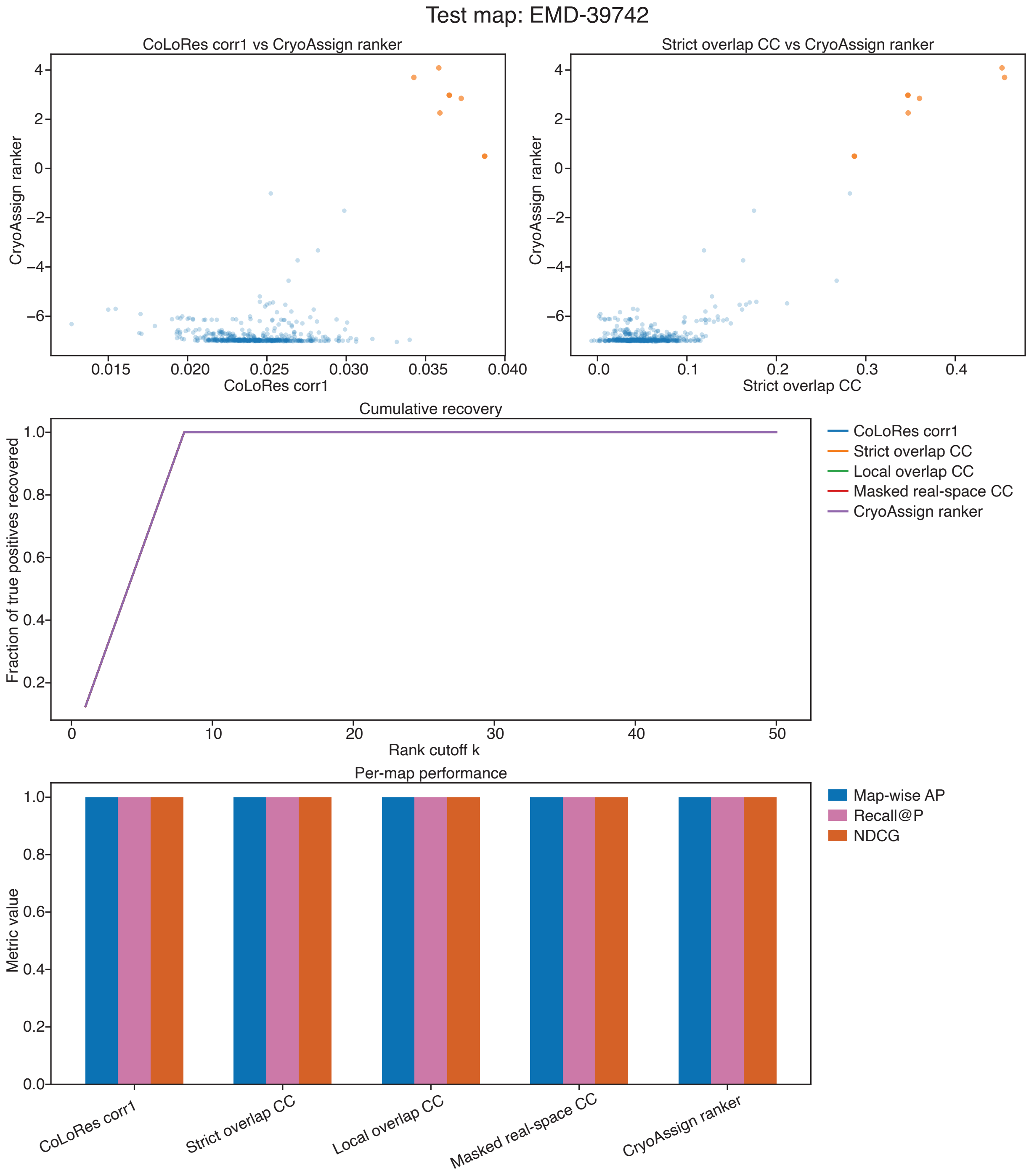

### Supplementary Figure 1i

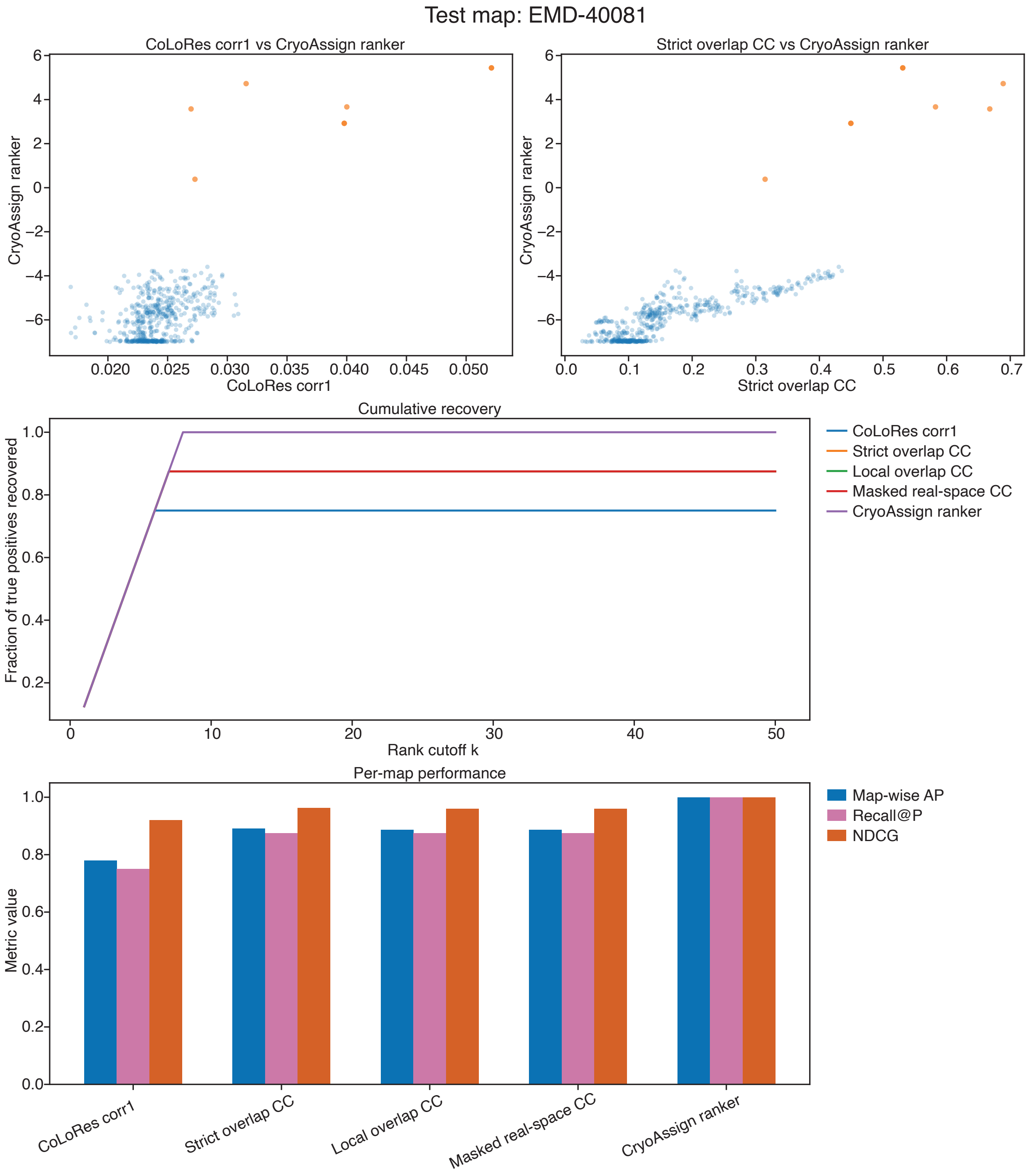

### Supplementary Figure 1j

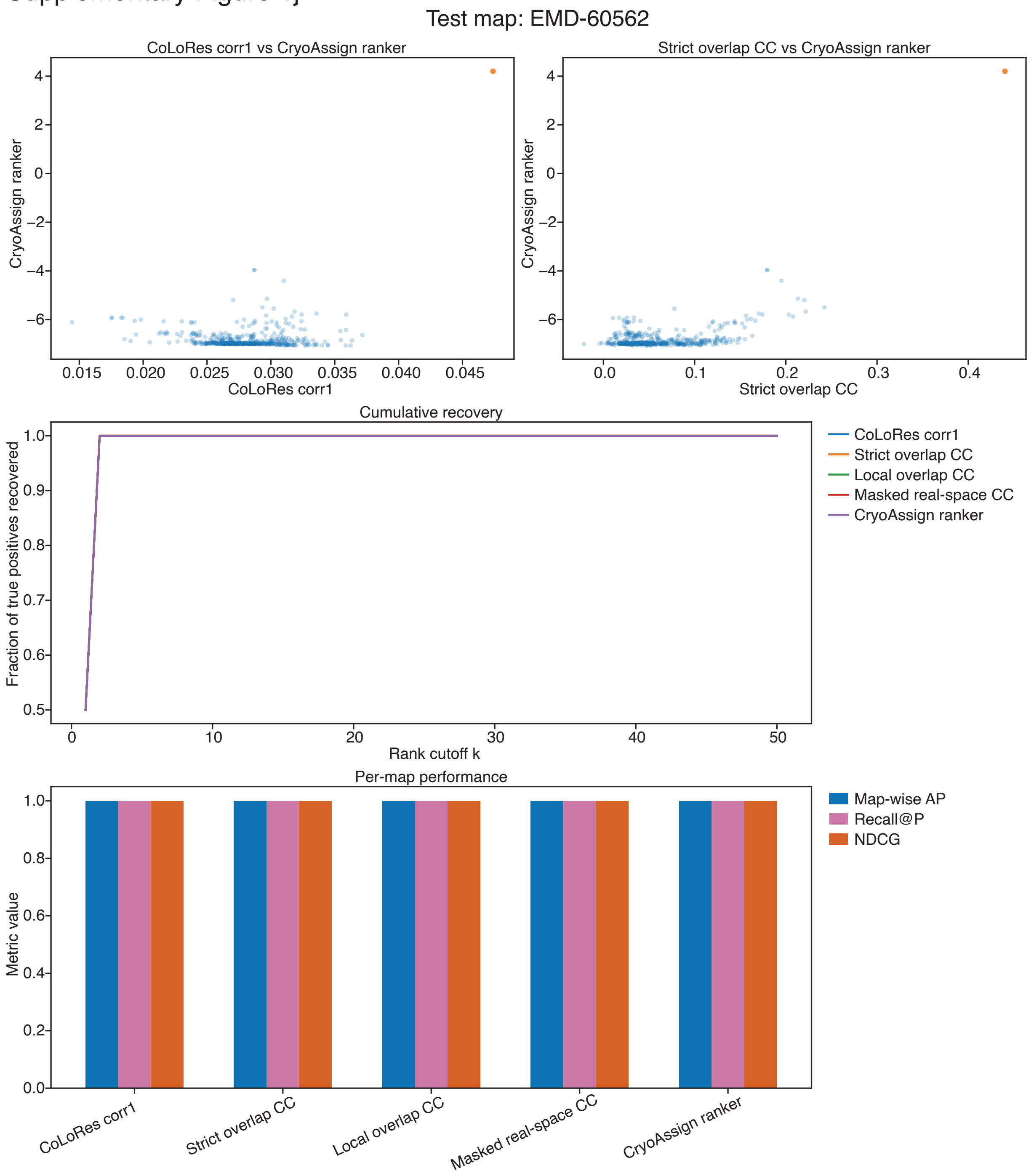

### Supplementary Figure 1k

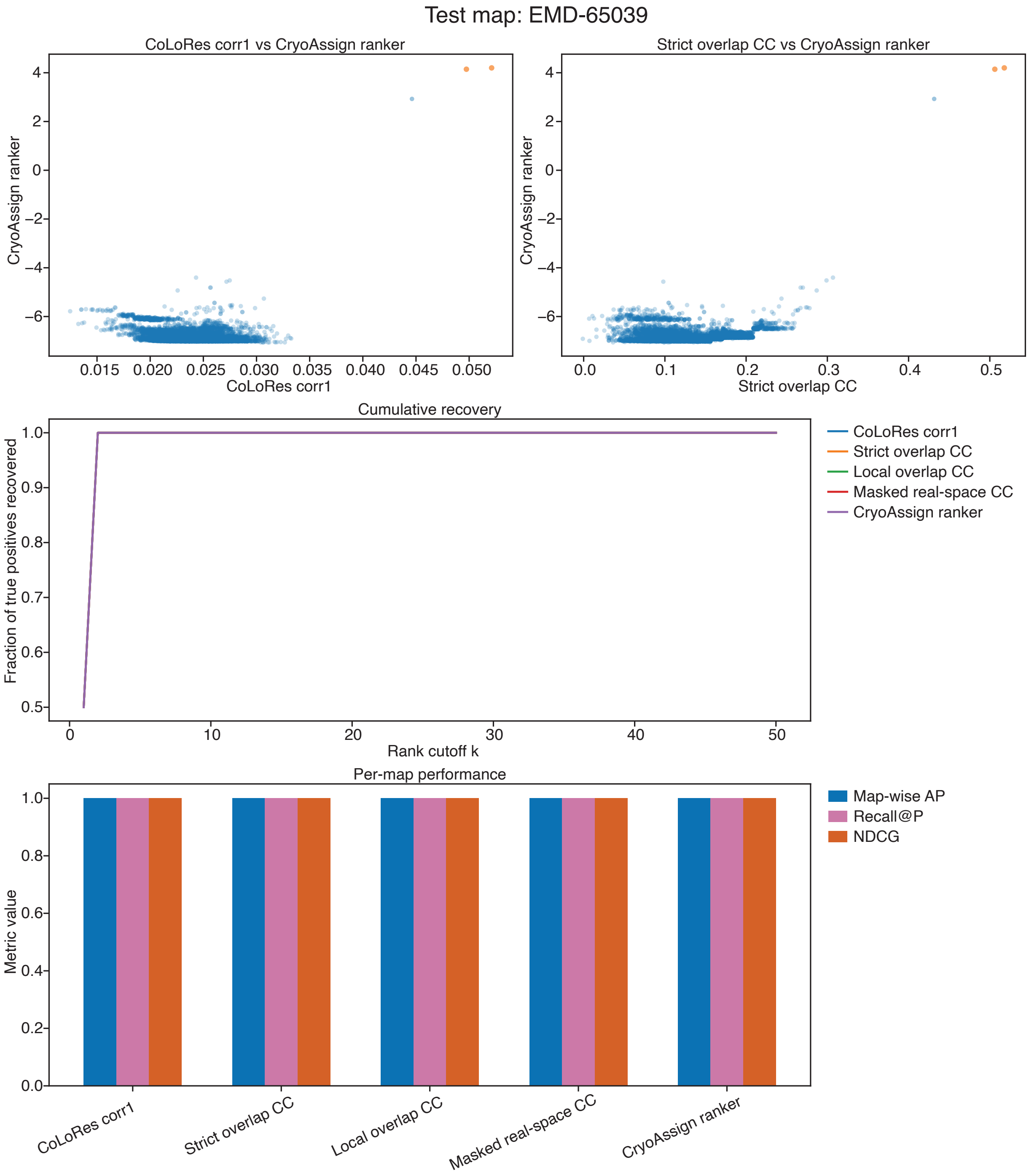

### Supplementary Figure 1l

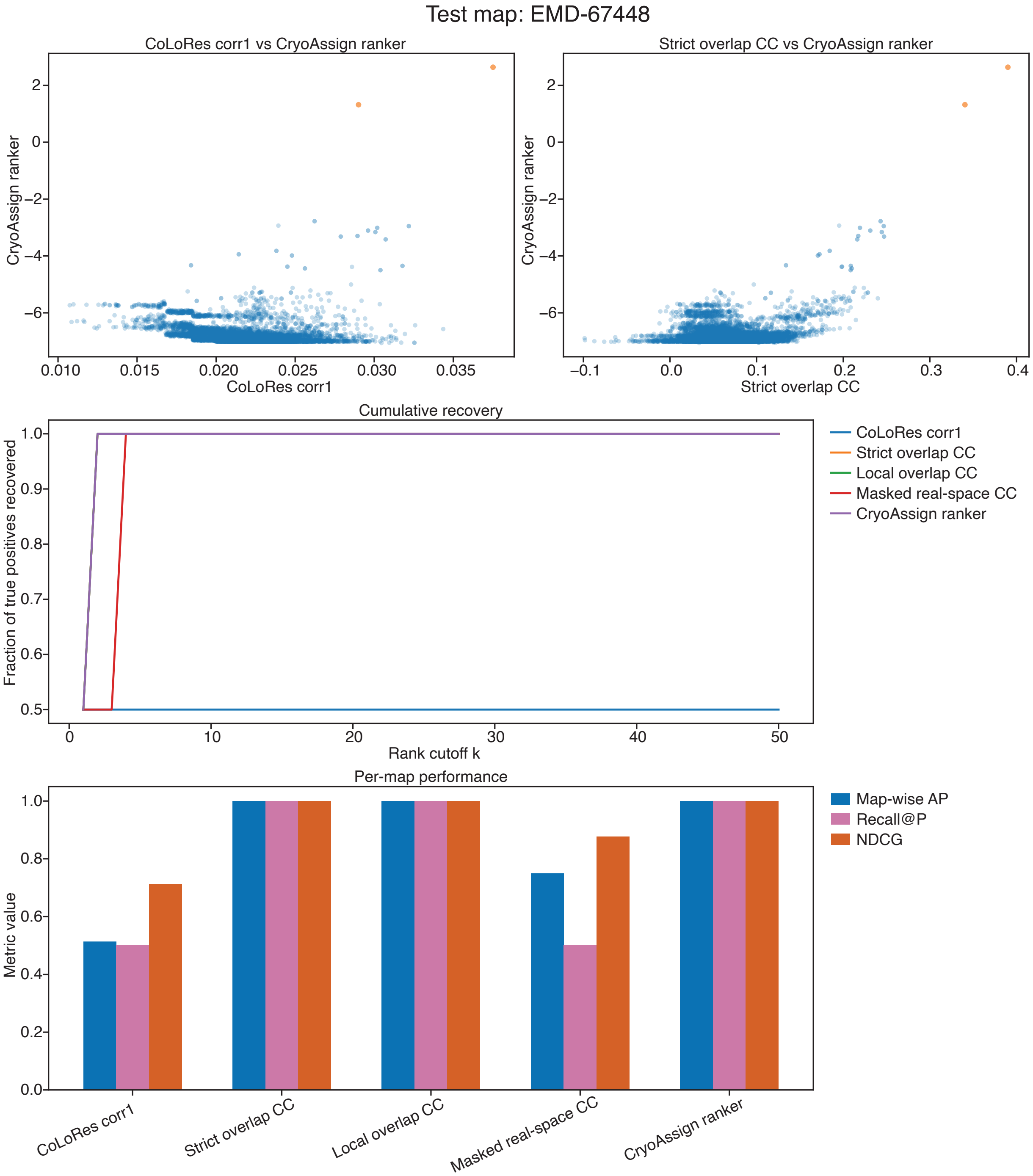

### Supplementary Figure 1m

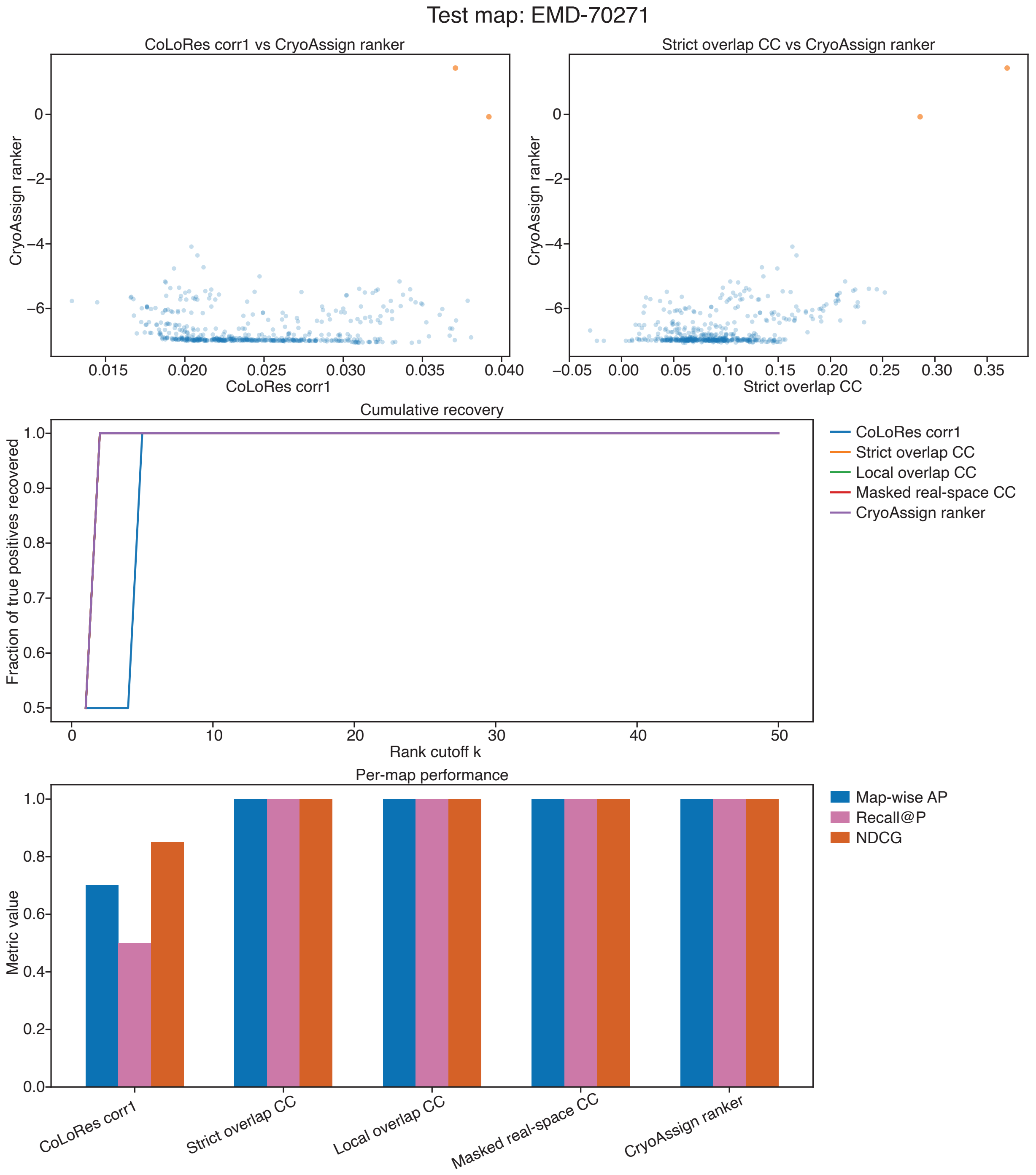

### Supplementary Figure 1n

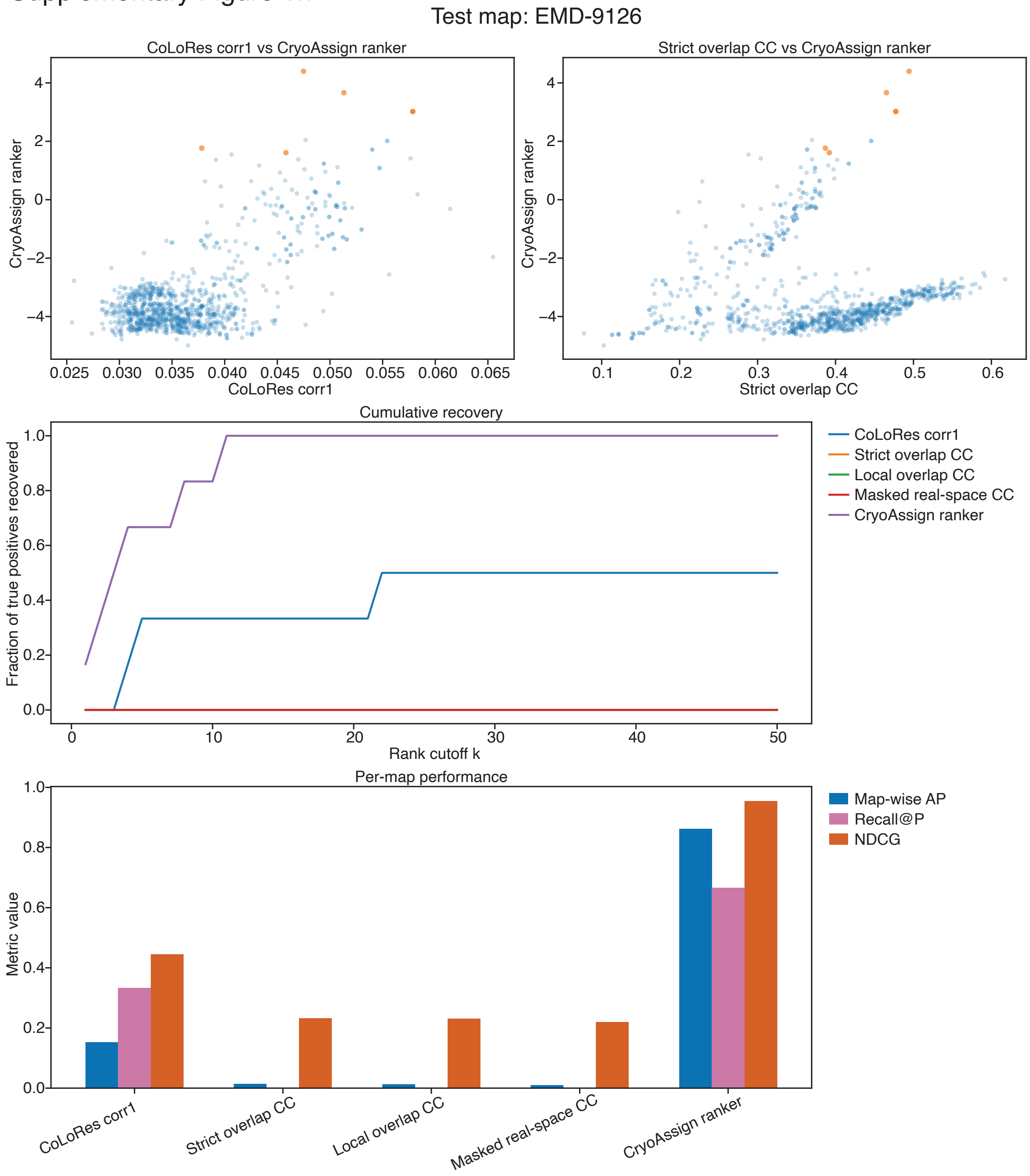

**Supplementary Figure 2 | Resolution-dependent changes in CryoAssign ranker scores and CoLoRes corr1 rankings.** CryoAssign was applied to EMD-9126, EMD-12661, EMD-50723, and EMD-65039 in their native forms and after progressive low-pass filtering, as described for Fig. 6. Panels follow the same map order as Fig. 6a–d. **a,c,e,g,** CryoAssign ranker scores for true-positive candidates and the 20 highest-scoring negative candidates at each filtering condition. The highest-scoring negative candidate at each condition is highlighted and connected across conditions by a dashed line. **b,d,f,h**, Ranks of the true-positive proteins obtained using the original CoLoRes corr1 score, displayed using the same grouping and axis conventions as in Fig. 6a–d.

Supplementary Figure 2

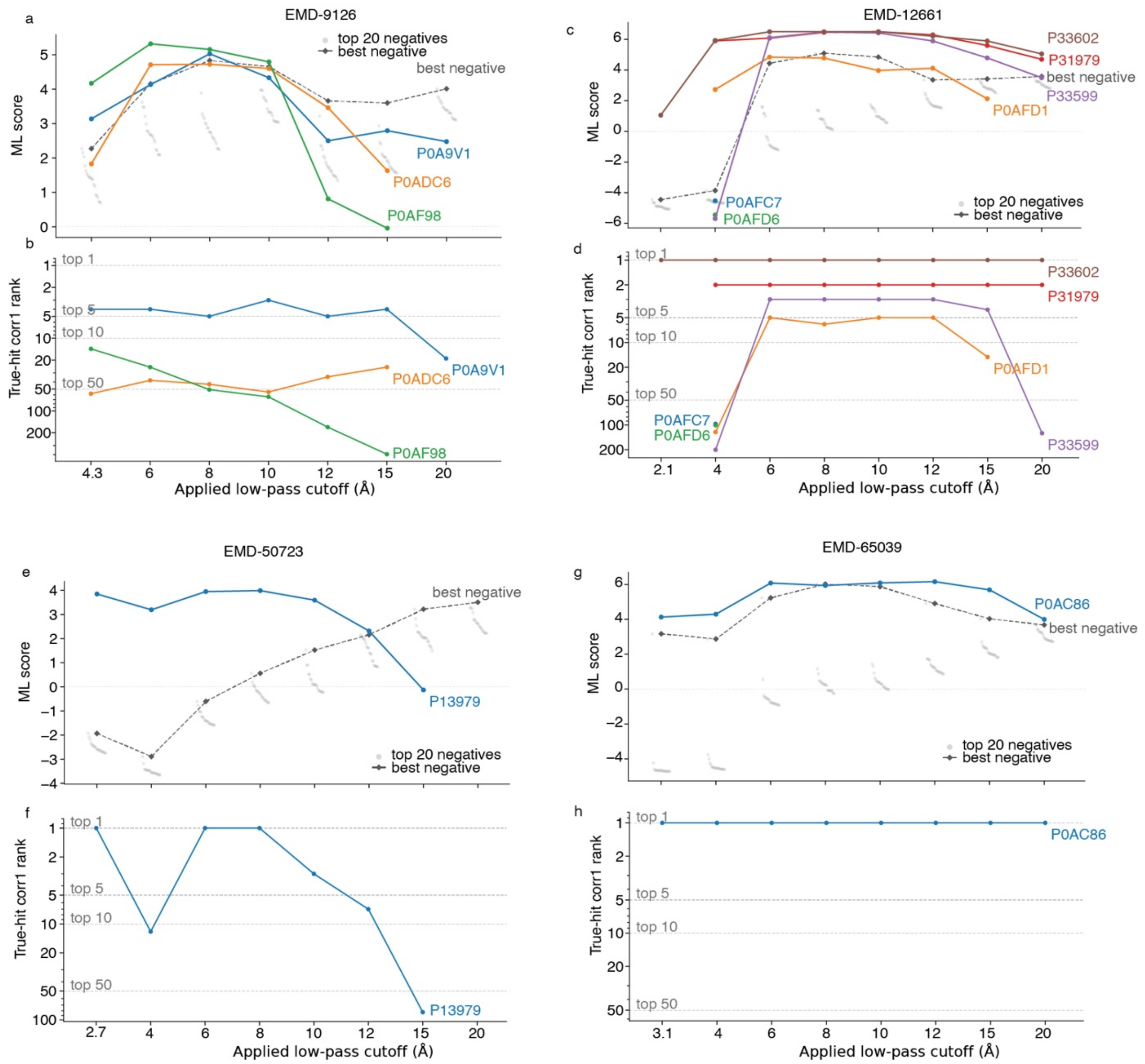
